# Microbiota-induced fatty acid synthesis facilitates intestinal infection and immune-mediated damage in *Drosophila*

**DOI:** 10.64898/2026.04.14.718535

**Authors:** Yi Yu, Kathirvel Alagesan, Dagmar Frahm, Emmanuelle Charpentier, Igor Iatsenko

**Affiliations:** Research group Genetics of host-microbe interactions, Max Planck Institute for Infection Biology, Charitéplatz 1, 10117 Berlin, Germany; Max Planck Unit for the Science of Pathogens, Charitéplatz 1, 10117 Berlin, Germany; Institute of Biology, Humboldt Universität-zu-Berlin, D-10115 Berlin, Germany

**Author notes:** **Corresponding author:** Igor Iatsenko, **Email:**.

**Keywords:** microbiota, host-microbe interactions, immunopathology, antimicrobial peptides, virulence, fatty acids, *Drosophila*, *Lactiplantibacillus plantarum*

## Abstract

The role of the microbiota in facilitating infection is increasingly recognized, though the underlying mechanisms remain under investigation. In this study, we demonstrate that the gut microbiome, particularly *Lactiplantibacillus plantarum*, promotes enteric infection in *Drosophila melanogaster* by reshaping host physiology and immune responses. Upon gut colonization, *L. plantarum* enhances fatty acid (FA) production in the gut. Reducing FA synthesis improves host survival, and pathogen mutants deficient in FA utilization exhibit reduced virulence. FAs support pathogen growth, increase virulence, and enhance resistance to antimicrobial effectors, ultimately facilitating pathogen persistence in the gut. Persisting pathogens consequently overstimulate the immune system, leading to excessive production of antimicrobial peptides (AMPs). Flies lacking AMPs, particularly *Metchnikowin* and *Attacin D* mutants, show better survival during infection, implicating AMPs in immunopathology. Our findings identify a mechanism whereby microbiota-induced release of host FAs stimulates pathogen virulence and persistence, ultimately driving AMP-mediated immunopathology.

## Introduction

Residing at the crossroads of host and environment, host-associated microbes or microbiota profoundly shape host-pathogen interactions and contribute to disease development ^1–3^. The most illustrative example of the importance of such interactions is colonization resistance - a concept of protection of the host from pathogens by commensal bacteria ^4^. While commensals’ protective functions against infections are well documented, with many underlying mechanisms described, an emerging paradigm from microbiome studies is that specific community compositions or particular microbiome members can increase host susceptibility to infection ^5,6^. Commensal signals and metabolites can be leveraged by gut pathogens to potentiate disease and regulate virulence gene expression ^1,7^. For instance, *Bacteroides thetaiotaomicron* liberates fucose and sialic acid from host glycans and produces abundant succinate, which *EHEC*, *S. Typhimurium*, and *C. difficile* sense to promote their expansion in the mammalian intestine ^8,9^. The other microbiota-produced metabolites, like acetate, butyrate, and taurocholate can regulate virulence genes in a number of enteric pathogens ^10–13^. Beyond these few exceptions, the molecular mechanisms through which commensals promote infection are not well defined. The complexity of animal microbiota makes it challenging to pinpoint which taxa or consortia drive beneficial or harmful interactions with a given pathogen. As most mammalian microbes remain unculturable and not amenable to genetic tools, functional testing of candidate interactions is challenging.

*Drosophila melanogaster* provides a tractable system to define how resident microbes influence susceptibility to enteric pathogens, leveraging powerful genetics and conserved innate immune defence ^14–16^. Fruit flies employ multiple strategies to defend against intestinal pathogens. The peritrophic matrix acts as a physical shield, protecting gut epithelial cells from microbial intrusion ^17,18^. Upon pathogen detection, inducible defenses—such as the synthesis of antimicrobial peptides (AMPs) and immune effectors like iron-sequestering proteins—are initiated in distinct gut regions ^19–21^. The Immune deficiency (Imd) pathway serves as the principal regulator of antimicrobial defenses in the midgut, whereas certain AMPs, including Drosomycin-like 2 (Drsl2) and Drsl3, are governed by the JAK-STAT signaling pathway ^19^. Reactive oxygen species (ROS) are also rapidly generated in the gut in response to bacterial exposure ^22–24^. Although the direct bactericidal effect of ROS is controversial ^25^, growing evidence suggests they also serve as signaling molecules that promote gut peristalsis and facilitate pathogen clearance through increased defecation ^26–29^. Intestinal cell damage further activates multiple signaling pathways, triggering stem cell proliferation, tissue repair, and resilience responses ^30–32^. The coordinated interplay between antimicrobial defenses and tissue repair processes is crucial for the fly’s survival during intestinal infection.

There are only a few known microbes that can overcome the intestinal defense mechanisms and establish a lethal infection in the *Drosophila* gut ^14^. *Pseudomonas entomophila* (*Pe*), a Gram-negative bacterium originally isolated from fruit flies, is one of the pathogens that can persist in the *Drosophila* gut following oral ingestion ^33^. Prominent features of *Pe* pathogenesis are irreversible toxin-mediated intestinal damage and translational inhibition, which prevent intestinal repair and immune responses ^20^. A pore-forming toxin, Monalysin, is a key *Pe* virulence factor implicated in gut damage, inhibition of intestinal peristalsis, and prevention of pathogen expulsion ^23,34,35^. *Pe* as a natural *Drosophila* pathogen allows the exploration of the crosstalk between infectious microbes and the commensal gut microbiota.

Flies are associated with low-diversity communities (2–30 species), generally dominated by the following species: *Lactiplantibacillus plantarum*, *Levilactobacillus brevis*, *Acetobacter pomorum*, *A. pasteurianus*, and *Enterococcus faecalis* ^36–38^. Such a low-diversity community of *Drosophila* microbiota, all members of which are cultivable and amenable to genetic manipulations, facilitates functional studies ^39^. Some *Drosophila* commensals can protect flies from *P. aeruginosa* or *Serratia marcescens* infections ^40^, but the mechanism underlying this protection has not been explored. Several studies suggest that the gut microbiota can defend the host against invasive microbes by acidifying the local environment ^41,42^. In particular, lactic acid produced by *L. plantarum* lowers gut pH, thereby restricting the growth of invasive pathogens ^42^. Although the examples demonstrating the detrimental impact of fly microbiota on infection outcomes are also emerging ^43–45^, it is not known how common such cases are among gut commensals and what determines the beneficial or detrimental role of a particular commensal. Given the well-known effect of microbiota on fly metabolism and the role of metabolism in *Drosophila* defence against *Pe* ^23,46,47^, we hypothesize that modulation of host metabolism by microbiota plays a role in response to infection.

Using *Drosophila-Pe* model, we demonstrated that gut microbiome facilitates pathogen infection in *Drosophila melanogaster* through reshaping the host physiology and immune response. The gut microbiome boosts the production of specific fatty acids that promote pathogen growth and virulence, thereby increasing pathogen loads. Pathogen persistence further induces the overproduction of several antimicrobial peptides, contributing to host mortality. This study provides direct evidence that the gut microbiome facilitates pathogen infection by modulating host metabolism and immune response, highlighting the potential of targeting specific metabolic and immune pathways to prevent infection.

## Results

### Microbiota facilitates *P. entomophila* infection

To determine how microbiota affects *Drosophila* susceptibility to intestinal infection with *Pe*, we generated axenic flies without microbiomes by hypochlorite dechorionation of embryos and scored the survival of these flies and conventional flies with microbiota after *Pe* infection (Figure 1A). Axenic flies demonstrated improved survival compared to infected conventional flies, indicating a detrimental effect of the microbiota during *Pe* infection (Figure 1B). To test whether specific bacteria are responsible for this detrimental effect, we isolated the bacterial species from our conventional flies by plating their homogenates on MRS and mannitol agar. Based on the sequencing of 16S ribosomal RNA gene, the isolated bacteria were identified as *L. plantarum* (hereafter *Lp^YY^*), *Sphingomonas leidyi*, *Sphingomonas* sp., *Variovorax* sp. We then generated gnotobiotic animals by reassociating axenic flies with each of the isolated microbes. Flies monoassociated with *Sphingomonas* and *Variovorax* exhibited mild but statistically significant (p=0.006 *Sphingomonas;* p=0.012 *Variovorax,* Table S1) decrease in survival during *Pe* infection as compared to germ-free controls (Figure S1A). Reassociation with *Lp^YY^* led to even a stronger decline in survival relative to axenic flies (Figure 1C). Flies colonized with another *Lp* strain, *Lp^WJL^*, or with *L. brevis* also showed increased susceptibility to *Pe*, indicating that the effect is not specific to *Lp^YY^* strain or to *L. plantarum* species (Figure 1C). Survival of flies colonized with *Acetobacter malorum* or *A. pomorum* - species typically found in flies^39^ was not significantly different (p>0.1) compared to germ-free controls (Figure S1B), suggesting that the tested acetic acid bacteria do not affect *Drosophila* susceptibility to *Pe*. We also assessed the microbial burden of *Pe* and found that while *Pe* load was similar in *Acetobacter*-colonized and axenic flies (Figure S1C), the presence of *Lp* or *Lb* in the mono-colonized flies resulted in significantly higher levels of *Pe* compared to axenic flies at 6h and 9h post-infection (Figure 1D). Together, these results demonstrate that the microbiome, particularly *Lp*, facilitates *Pe* persistence in the gut, which is consistent with the decreased survival observed in *Lp*-colonized flies.

**Figure 1.**
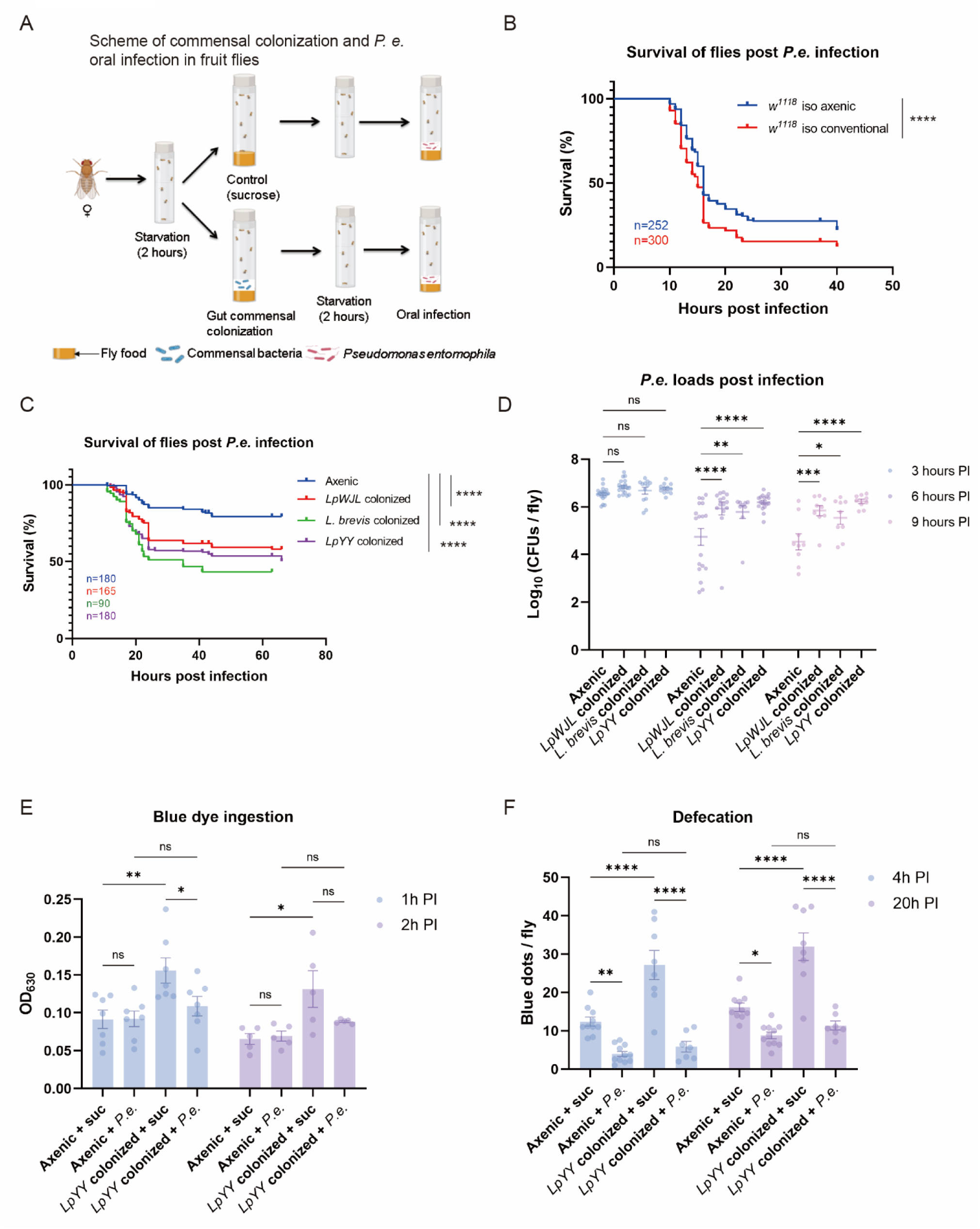
Microbiota facilitates *P. entomophila* infection. (A) Scheme showing the experimental procedure for pathogen oral infection (without the commensal colonization step) and gut commensal bacteria colonization following pathogen oral infection. (B) Survival curves of conventional and axenic *w^1118^* iso flies upon *P. entomophila* infection. The accumulated survival graph is shown (*n* = 4). (C) Survival curves of *Lactiplantibacillus*-colonized axenic *w^1118^* iso flies upon *P. entomophila* infection. Three *Lactiplantibacillus* species were used: *L. plantarum WJL* (*LpWJL*), *L. plantarumYY* (*LpYY*) and *L. brevis*. The accumulated survival graph is shown (*n* = 2-4). (D) Pathogen loads (CFUs) at 3, 6, and, 9 hours post-exposure to *P. entomophila* in axenic and *Lactiplantibacillus*-colonized *w^1118^* iso flies. Each dot represents a pool of five flies taken from different infection vials (*n* ≥ 9, pooled from at least 3 experiments). Bars show mean ± SEM. (E) Ingestion of blue dye mixed with *P. entomophila* during 1 and 2 hours of exposure in *w^1118^*iso flies. Each dot represents a pool of five flies taken from different infection vials (*n* ≥ 5, pooled from at least 2 experiments). Bars show mean ± SEM. (F) The defecation rate of axenic and *Lactiplantibacillus*-colonized *w^1118^* iso flies was measured 4 and 20 hours after exposure to blue-dyed sucrose (control) or *P. entomophila*. Each dot indicates a pool of 10 flies taken from different infection vials (*n* ≥ 7, pooled from at least 2 experiments). Bars show mean ± SEM. Data analyzed with stratified Cox hazard ratios (B and C), two-way ANOVA plus Sidak’s test (D), and Tukey’s test (E and F). ns, not significant; \**p* < 0.05, \*\**p* < 0.01, \*\*\**p* < 0.001, \*\*\*\**p* < 0.0001. For detailed sample sizes and statistical analyses, see Table S1.

We then focused on dissecting the mechanisms underlying the detrimental effect of *Lp* during infection with *Pe*. We hypothesized that *Lp* might alter fly feeding behaviour, leading to an increased pathogen ingestion. To assess the amount of solution that flies ingest, we added blue dye to sucrose or *Pe* suspension and quantified the intensity of blue colour in fly homogenates. We found that *Lp*-colonized flies ingest significantly more of sucrose (control) solution as compared to axenic flies (Figure 1E). However, the pathogen uptake was comparable between axenic and *Lp*-associated flies at 1h and 2h post infection (Figure 1E). Hence, axenic and *Lp*-colonized flies ingest a similar pathogen amount, which is also reflected in a similar pathogen load at 3h after infection (Figure 1D). Given that we previously found that *Pe* inhibits defecation and pathogen clearance ^23^, we investigated whether *Lp* could exacerbate this process and facilitate *Pe* persistence. However, we found that *Lp*-associated flies showed the same defecation frequency as axenic controls 4h and 20h post *Pe* infection (Figure 1F), suggesting that *Pe* persistence in *Lp*-colonized flies is not due to reduced pathogen clearance via defecation. Also, we did not detect *Pe* or *Lp* in the hemolymph of all treatment groups (Figure S1D), excluding systemic infection by a pathogen or commensal as a cause of death in *Lp* colonized flies.

### Exaggerated immune activation upon *Pe* infection in *Lp*-colonized flies

To investigate how *Lp* shapes the host response to *Pe* infection and identify affected host processes that might contribute to increased susceptibility of gnotobiotic flies, we investigated the intestinal response of axenic and *Lp*-colonized flies to *Pe* infection using RNA-seq. Uninfected axenic and *Lp*-colonized flies served as additional controls allowing us to determine gut response to *Lp* colonization or *Pe* infection. As anticipated from previous studies ^48,49^, *Lp* colonization (*Lp* colonized vs. UC (axenic)) had a rather mild effect on changes in gene expression. We found 112 genes up- and 36 down-regulated in *Lp* colonized flies (Figure 2A, Table S2). Negative regulators of Imd pathway (*pirk, PGRP-SC1a, PGRP-SC1b*) were induced by *Lp* colonization, likely preventing the AMP response, as no AMPs were induced in *Lp* colonized flies. Genes involved in lipid and protein metabolism were upregulated upon *Lp* colonization. *Pe* infection (*Pe* vs UC) triggered broader changes in gene expression with 2808 genes up- and 1583 down-regulated (Figure 2B, Table S2). As expected, immune (AMPs: *DptA/B, AttC, Mtk*), epithelial repair (*upd3, Socs36E, pvf1*) stress response, and detoxification (*Tots, Gsts, Hsps*) genes were induced, while metabolism genes were repressed by *Pe* infection. *Pe* infection of *Lp* colonized flies (*LpPe* vs UC) induced a similar response to *Pe* infection alone with immune, stress response, and epithelial renewal genes being up-regulated (Figure 2C, Table S2). This similarity is also reflected in the number of overlapping genes: 899 genes were common to the *Pe* and *LpPe* treatment groups (Figure 2D, Table S3), indicating that these genes were differentially regulated by *Pe* independently of *Lp* presence. We identified 127 genes that were uniquely differentially expressed in response to *Pe* infection in *Lp*-associated flies (Figure 2D, Table S3). Besides several immune genes, such as *Daisho2, BomBc1, Tep3, Hayan* we did not detect any particular expression pattern that could be linked to increased susceptibility to infection (Table S3). Hence, we decided to check genes that were differentially expressed between axenic and *Lp*-colonized flies in response to *Pe* infection (*LpPe vs Pe*). There were 152 genes up- and 281 down-regulated in *Lp*-colonized flies (Figure 2E, Table S2). Gene Ontology (GO) analysis of *LpPe*-upregulated genes identified several GO categories related to defense and immune response, indicating an enrichment of immune response genes in *LpPe* dataset (Figure 2F). The heat map of the selected immune genes further confirms their higher expression level in *LpPe* flies as compared to the other treatment groups (Figure 2G). Together, these results illustrate a stronger immune response of *Lp*-colonized flies to *Pe* infection as compared to axenic controls.

**Figure 2.**
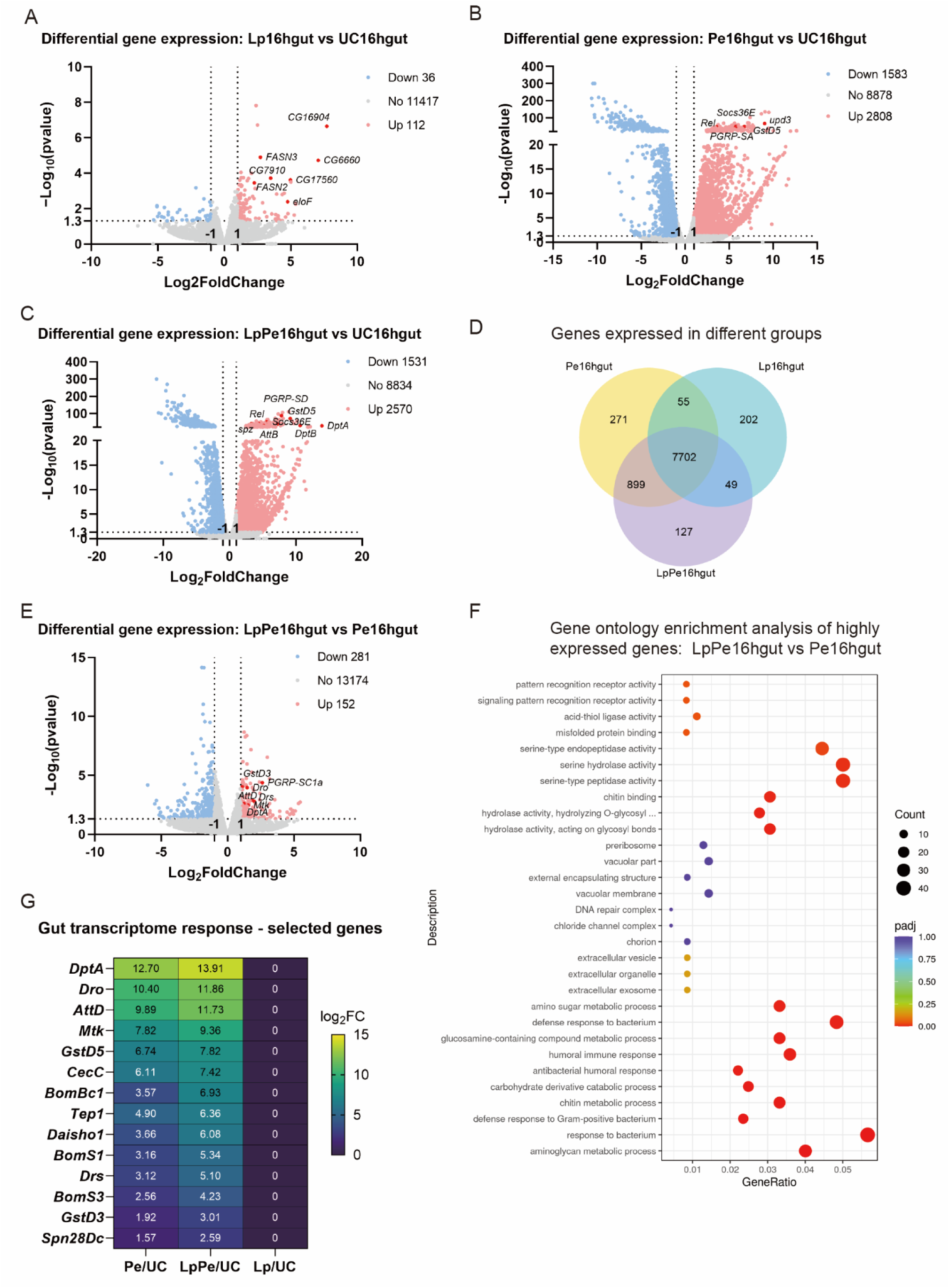
Exaggerated immune activation upon *Pe* infection in *Lp*-colonized flies. (A) Volcano plot showing changes in gene expression in the guts of *Lactiplantibacillus*-colonized *w^1118^* iso flies with a 16-hour sucrose treatment (Lp16hgut) versus unchallenged flies with a 16-hour sucrose treatment (UC16hgut) (*n* = 3). Numbers indicate significantly regulated genes (*p* < 0.05 and |Log_2_FoldChange| ≥ 1). (B) Volcano plot showing gene expression changes in guts of axenic *w^1118^* iso flies upon a 16-hour *P. entomophila* infection (Pe16hgut) versus unchallenged flies with a 16-hour sucrose treatment (UC16hgut) (*n* = 3). Numbers indicate significantly regulated genes (*p* < 0.05 and |Log_2_FoldChange| ≥ 1). (C) Volcano plot showing gene expression changes in the guts of *Lactiplantibacillus*-colonized *w^1118^* iso flies upon a 16-hour *P. entomophila* infection (LpPe16hgut) versus unchallenged flies with a 16-hour sucrose treatment (UC16hgut) (*n* = 3). Numbers indicate significantly regulated genes (*p* < 0.05 and |Log_2_FoldChange| ≥ 1). (D) Venn diagram plot showing overlapping and unique genes expressed in *Lactiplantibacillus*-colonized flies alone (Lp16hgut), axenic flies upon 16-hour *P. entomophila* infection (Pe16hgut), and *Lactiplantibacillus*-colonized flies upon 16-hour *P. entomophila* infection (LpPe16hgut). (E) Volcano plot showing gene expression changes in the guts of *Lactiplantibacillus*-colonized *w^1118^* iso flies upon a 16-hour *P. entomophila* infection (LpPe16hgut) versus axenic *w^1118^* iso flies upon a 16-hour *P. entomophila* infection (Pe16hgut) (*n* = 3). Numbers indicate significantly regulated genes (*p* < 0.05 and |Log_2_FoldChange| ≥ 1). (F) Dot map showing Gene Ontology enrichment analysis of highly expressed genes in the guts of *Lactiplantibacillus*-colonized *w^1118^* iso flies upon a 16-hour *P. entomophila* infection (LpPe16hgut) and axenic *w^1118^* iso flies upon a 16-hour *P. entomophila* infection (Pe16hgut) (*n* = 3). (G) Heatmap showing selected significantly upregulated genes in the guts of *Lactiplantibacillus*-colonized (Lp16hgut) flies, the guts of axenic *w^1118^* iso flies upon a 16-hour *P. entomophila* infection (Pe16hgut), and the guts of *Lactiplantibacillus* colonized *w^1118^*iso flies upon a 16-hour *P. entomophila* infection (LpPe16hgut) versus unchallenged flies with a 16-hour sucrose treatment (UC16hgut). Log_2_FC value indicates the average of 3 experiments.

### Overexpression of antimicrobial peptides decreases the survival of *Pe*-infected flies

We hypothesized that excessive immune response in *Lp*-colonized flies could contribute to their increased susceptibility to *Pe* infection. To test this hypothesis, we investigated whether the effect of *Lp* is present in immunodeficient flies. Specifically, we used *Relish* mutants, which are deficient for the Imd pathway – a key humoral immune response pathway in the gut ^15^. We observed comparable survival of axenic and *Lp*-colonized *Relish* mutant flies after *Pe* infection (Figure 3A). We further confirmed these findings with gut-specific knockdown of *Relish* by RNAi (Figure S2A), indicating that a functional Imd pathway is necessary for *Lp* to promote *Pe* infection. To test whether the involvement of Imd pathway is linked to the production of immune effectors, we tested *ΔAMP* mutant flies lacking ten AMP genes, including 4 Attacins, 2 Diptericins, Drosocin, Drosomycin, Metchnikowin and Defensin ^50^. Unexpectedly, *ΔAMP* mutant flies both axenic and *Lp*-colonized showed increased survival rates during *Pe* infection as compared to the respective wild-type controls (Figure 3B). *Lp*-colonized *ΔAMP* mutant flies demonstrated higher survival than axenic controls—a phenotype opposite to that of wild-type flies (Figure 3B). Consistent with survival, there was a lower *Pe* burden in *Lp*-colonized *ΔAMP* mutant flies as compared to wild-type controls (Figure 3C). In contrast to wild-type flies that had higher *Pe* load in *Lp*-colonized *versus* axenic state, *Relish* mutants contained equally high pathogen load under axenic and *Lp*-colonized conditions (Figure 3C), consistent with equal susceptibility of these flies to *Pe* (Figure 3A).

**Figure 3.**
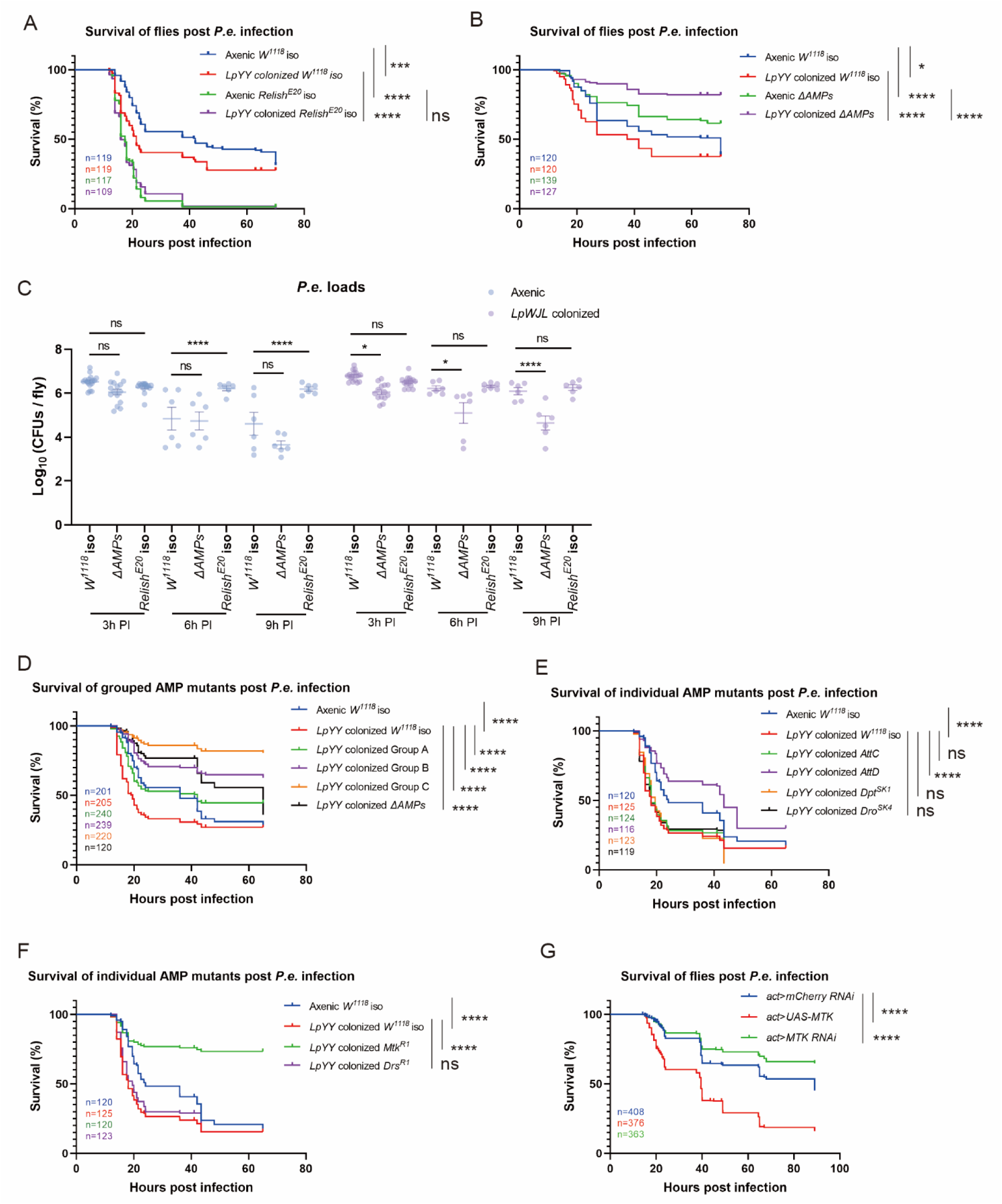
Overexpression of antimicrobial peptides decreases survival of *Pe*-infected flies. (A) Survival curves of *Lactiplantibacillus*-colonized axenic *w^1118^* iso flies and *Relish^E20^-*mutated flies (*Relish^E20^ iso*) upon *P. entomophila* infection. The accumulated survival graph is shown (*n* = 3). (B) Survival curves of *Lactiplantibacillus*-colonized axenic *w^1118^* iso flies and flies mutated in 10 antimicrobial peptide genes (*ΔAMPs*) upon *P. entomophila* infection. The accumulated survival graph is shown (*n* = 3). (C) Pathogen loads (CFUs) at 3-, 6-, and 9-hours post-exposure to *P. entomophila* in axenic and *Lactiplantibacillus*-colonized *w^1118^* iso, *Relish^E20^ iso* and *ΔAMPs* flies. Each dot indicates a pool of five flies taken from different infection vials (*n* ≥ 6, pooled from at least 2 experiments). Bars show mean ± SEM. (D) Survival curves of *Lactiplantibacillus*-colonized axenic *w^1118^* iso flies and flies mutated in antimicrobial peptide genes (Group A: *defensin;* Group B: *drosocin*, *diptericin A-B*, and *Attacin A-D;* Group C: *drosomycin* and *metchnikowin*) upon *P. entomophila* infection. The accumulated survival graph is shown (*n* = 3-5). (E) Survival curves of *Lactiplantibacillus*-colonized axenic *w^1118^* iso flies and flies mutated in Group B antimicrobial peptide genes (Group B: *drosocin*, *Dro^SK4^*; *diptericin, Dpt^SK1^; AttacinC, AttC and Attacin D*, *AttD*) upon *P. entomophila* infection. The accumulated survival graph is shown (*n* = 3-5). (F) Survival curves of *Lactiplantibacillus*-colonized axenic *w^1118^* iso flies and flies mutated in Group C antimicrobial peptide genes (Group C: *drosomycin, Drs^R1^*and *metchnikowin, Mtk^R1^*) upon *P. entomophila* infection. Accumulated survival graph is shown (*n* = 3). (G) Survival curves of *conventional* flies overexpressing the antimicrobial peptide *metchnikowin (act>UAS-MTK)* or downregulating with *metchnikowin (act>MTK RNAi)*. mCherry RNAi was used as a control. The accumulated survival graph is shown (*n* = 8). Data analyzed with stratified Cox hazard ratios (A, B, D, E, F, and G) and one-way ANOVA plus Sidak’s test (C). ns, not significant; \**p* < 0.05, \*\**p* < 0.01, \*\*\**p* < 0.001, \*\*\*\**p* < 0.0001. For detailed sample sizes and statistical analyses, see Table S1.

Next, we attempted to identify a specific AMP or group of AMPs that are detrimental during *Pe* infection. For this, we applied AMP-groups approach ^50^ by testing the survival of *Lp*-colonized mutants that lack specific groups of AMPs: Group A-flies lacking *Defensin (Def)*; Group B- flies lacking *Drosocin (Dro)*, *Diptericins (Dpt)* and *Attacins (Att)*; Group C- flies lacking *Metchnikowin (Mtk)* and *Drosomycin (Drs)*. While all three groups showed increased survivals, group C survived *Pe* infection even better than *ΔAMP* mutant (Figure 3D). Next, we tested individual mutants from group A and B and *ΔCec^A-C^*mutant lacking the four *cecropin* genes, *CecA1, CecA2, CecB*, and *CecC* ^51^. Most of the mutants, including *ΔCec^A-C^*, *AttC, Dro^SK4^, Dpt^SK1^* survived to the same degree as wild-type (Figure 3E, Figure S2B). *AttD* mutant, however, survived better than wild-type (Figure 3E). We also tested individual *Mtk* and *Drs* mutants that together constitute group C flies and found that *Mtk* but not *Drs* mutants survived *Pe* infection better than wild-type flies (Figure 3F). To reinforce the phenotype of *Mtk* mutant, we performed knockdown and overexpression experiments. *Mtk* RNAi flies survived *Pe* infection significantly better than control flies, whereas *Mtk* overexpression significantly decreased the survival rate of flies (Figure 3G). Overall, these results support a detrimental effect of Mtk and AttD on *Drosophila* survival during *Pe* infection. Given that *Pe* infection strongly induces *Mtk* and *AttD* expression in the gut (Figure 2G)—particularly in *Lp*-colonized flies—overactivation of these AMPs in flies likely contributes to their heightened susceptibility to *Pe* infection.

### *Lp* triggers the expression of FA genes and increases FA amounts in the gut

Next, we investigated whether direct interactions between *Pe* and *Lp* or modulation of host physiology by *Lp* facilitate *Pe* persistence in the guts of *Lp*-colonized flies. We co-cultured *Lp* and *Pe* in BHI media and found that *Pe* reached the same quantity in co-culture as in mono-culture, suggesting that *Lp* does not affect the growth of *Pe* (Figure S3A). Hence, we hypothesized that *Lp* alters gut physiology in a manner that supports *Pe* pathogenesis. In such a case, we would anticipate that the effect of *Lp* would be time-dependent as changes that it triggers unlikely to occur instantly. Indeed, we found that while flies pre-colonized with *Lp* for 24h (our standard protocol) exhibited increased susceptibility to *Pe* in comparison with axenic flies (Figure 4A), flies pre-colonized for only 4h were as susceptible as axenic controls (Figure S3B). Additionally, flies that received *Lp* simultaneously with *Pe* survived even better than axenic controls (Figure 4A). Such time-dependent effect of *Lp* is consistent with the hypothesis that *Lp* facilitates *Pe* infection by altering the host physiology. To find how exactly *Lp* shapes the host physiology, we checked our RNAseq results for the processes affected by *Lp* colonization in flies. Gene Ontology analysis of genes upregulated in *Lp*-colonized flies identified two major GO categories: endopeptidase activity and fatty acid synthesis, suggesting that *Lp* induces proteolytic enzymes, as previously reported in larvae ^52^, and fatty acid synthesis (Figure 4B). To test whether enhanced intestinal proteolytic activity contributes to *Lp*-mediated infection facilitation, we pre-fed flies with different concentrations of protease inhibitor prior to infection. Under these conditions, *Lp*-colonized flies were still more susceptible to *Pe* infection in comparison with axenic controls, suggesting that higher protease activity in *Lp*-colonized guts does not play a role in *Pe* infection (Figure S3C). To address the contribution of fatty acid synthesis, we first tested whether *Lp*-colonized flies in comparison with axenic controls contain more fatty acids (FAs) as a consequence of increased expression of fatty acid synthesis genes. Indeed, we detected a significantly higher amount of free FAs in the guts of *Lp*-colonized flies (Figure 4C). Importantly, flies colonized with *A. malorum* – a microbe that does not promote *Pe* infection, contained similar to axenic flies amounts of FAs (Figure S3D), suggesting that the effect on FAs synthesis is not general to any microbe. Next, we performed LC-ESI-MS/MS analysis of FAs in axenic and *Lp*-colonized flies to identify and quantify specific FAs. Overall, we detected FAs with high (>100 pmol/mg) and low abundance (<100 pmol/mg) (Figure 4D, 4E). Most of the detected FAs were present in similar quantities. However, *Lp*-colonized samples had a significantly higher amount of lauric acid (FA 12:0;0), myristoleic acid (FA 14:1;0), palmitoleic acid (FA 16:1;0), oleic acid (FA 18:1;0). To test whether elevated amount of FAs in *Lp*-colonized flies contributes to their increased susceptibility to *Pe* infection, we took two approaches. First, we fed flies palmitoleic acid (FA 16:1;0)—the most abundant FA in our samples and elevated in *Lp*-colonized flies—for 8 days to achieve saturation. Such flies exhibited increased susceptibility to *Pe* infection as compared to flies maintained on a control food (Figure 4F). Feeding flies with oleic (FA 18:1;0) or myristoleic (FA 14:1;0) acids - both of which are more abundant in *Lp*-colonized flies – had no significant effect on survival (Figure 4F), suggesting that only specific FAs correlate with increased susceptibility to *Pe* infection. Second, we genetically manipulated flies to reduce the synthesis on FAs by knocking down Acetyl-CoA carboxylase (*acc*) gene which encodes a key enzyme in FAs synthesis ^53^. We found that *acc* RNAi flies survived *Pe* infection better then control flies under *Lp* colonized conditions (Figure 4G). We further reinforced this result with knockdown of Fatty acid synthase 2 (<u>FASN2</u>) – a gene that was upregulated in *Lp* colonized flies and is needed for the synthesis of long chain fatty acids ^53^ (Figure S3E). These results suggest that *Lp* facilitates *Pe* infection by increasing the production of of FAs by the host.

**Figure 4.**
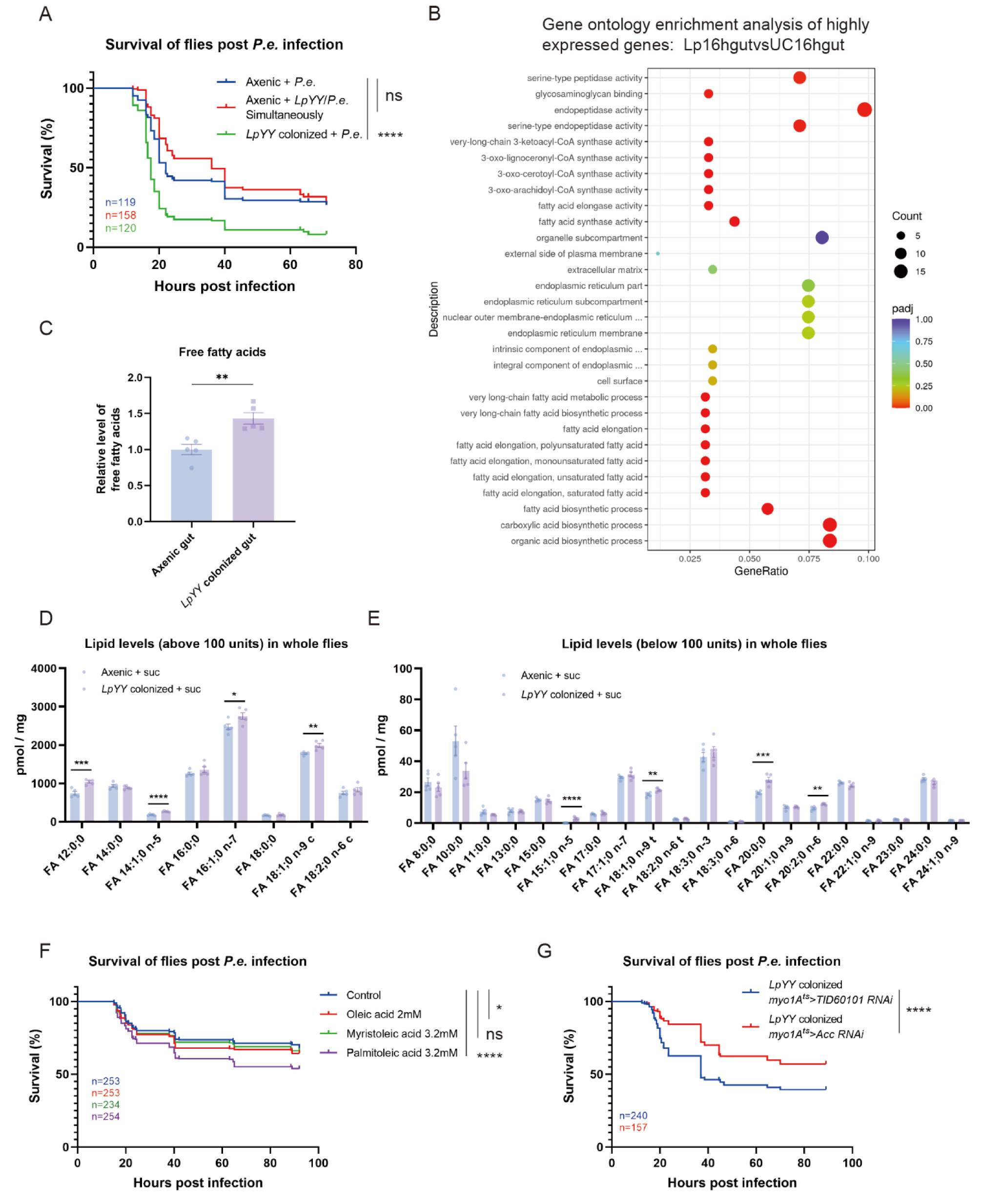
*Lp* triggers the expression of fatty acid (FA) genes and increases FA amounts in the gut. (A) Survival curves of *Lactiplantibacillus*-colonized axenic *w^1118^* iso flies upon *P. entomophila* infection and flies treated with a mixture of *Lactiplantibacillus* and *P. entomophila* simultaneously. The accumulated survival graph is shown (*n* = 3). (B) Dot map showing gene ontology enrichment analysis of highly expressed genes in the guts of *Lactiplantibacillus*-colonized *w^1118^* iso flies after 16 hours of *P. entomophila* infection (LpPe16hgut) versus unchallenged axenic *w^1118^* iso flies (UC16hgut) (*n* = 3). (C) Free fatty acid measurements from the guts of *Lactiplantibacillus*-colonized axenic *w^1118^*iso flies and axenic flies. The relative level of free fatty acids was normalized by setting the value from the axenic gut to 1. Each dot indicates a pool of 20 fly guts taken from different infection vials (*n* = 5, pooled from 2 experiments). Bars show mean ± SEM. (D) Levels of lipids above 100 pmol/mg of extracted material from whole flies colonized with *Lactiplantibacillus* and axenic flies. Each dot indicates a pool of 20 flies taken from different infection vials (*n* = 5, pooled from 2 experiments). Bars show mean ± SEM. (E) Levels of lipids below 100 pmol/mg of extracted material from whole flies. Samples were analyzed the same as in (D). (F) Survival curves of axenic *w^1118^* iso flies fed with fatty acids for 8 day and exposed to *P. entomophila*. The accumulated survival graph is shown (*n* = 5). (G) Survival curves of *Lactiplantibacillus-*colonized *w^1118^* iso flies with the gut specific downregulation of acetyl CoA carboxylase (*myo1A^ts^>Acc RNAi*) and using the TID60101 RNAi line as a control (*myo1A^ts^>* TID60101 RNAi). The accumulated survival graph is shown (*n* = 4). Data analyzed with stratified Cox hazard ratios (A, F, and G) and an unpaired t-test (C, D and E). ns, not significant; \**p* < 0.05, \*\**p* < 0.01, \*\*\**p* < 0.001, \*\*\*\**p* < 0.0001. For detailed sample sizes and statistical analyses, see Table S1.

### FAs increase *Pe* virulence

To understand the role of FAs in *Pe* infection, we investigated their impact on *Pe*. First, we tested the effect of FAs on *Pe* growth in M9 minimal media supplemented with any of the four FAs that were significantly more abundant in *Lp*-colonized flies. We found that *Pe* can grow and hence utilize any of the four tested FAs as a sole carbon and energy source (Figure 5A). Additionally, we investigated whether FAs can protect *Pe* against AMPs, using cationic antimicrobial polymyxin B that mimics the action of some *Drosophila* AMPs. Supplementation of LB media with palmitoleic acid enabled *Pe* to grow in the presence of 8 µg/ml of polymyxin B, while the non-supplemented control grew only at 1 µg/ml (Figure 5B). Although to a lesser extent, myristoleic acid but not lauric acid also significantly increased *Pe* ability to grow in the presence of polymyxin B (Figure S4A, S4B). A similar effect of FAs was observed in M9 minimal media (Figure S4C-E). Hence, FAs promote growth and the ability to survive AMPs. To test whether FAs can promote *Pe* virulence, we pre-incubated *Pe* for 3h with specific FAs prior to conducting systemic infection by pricking. We used systemic infection here as it is a more controlled and sensitive method compared to oral infection, which also requires a high concentration of infection dose. Flies that were infected with *Pe* pre-incubated with palmitoleic acid had significantly reduced survival compared to control flies infected with LB pre-incubated *Pe* (Figure 5C). Pre-incubation with lauric or myristoleic acid had no significant impact on *Pe* ability to kill flies, suggesting that only specific FAs, like palmitoleic acid, can increase *Pe* virulence (Figure S4F).

**Figure 5.**
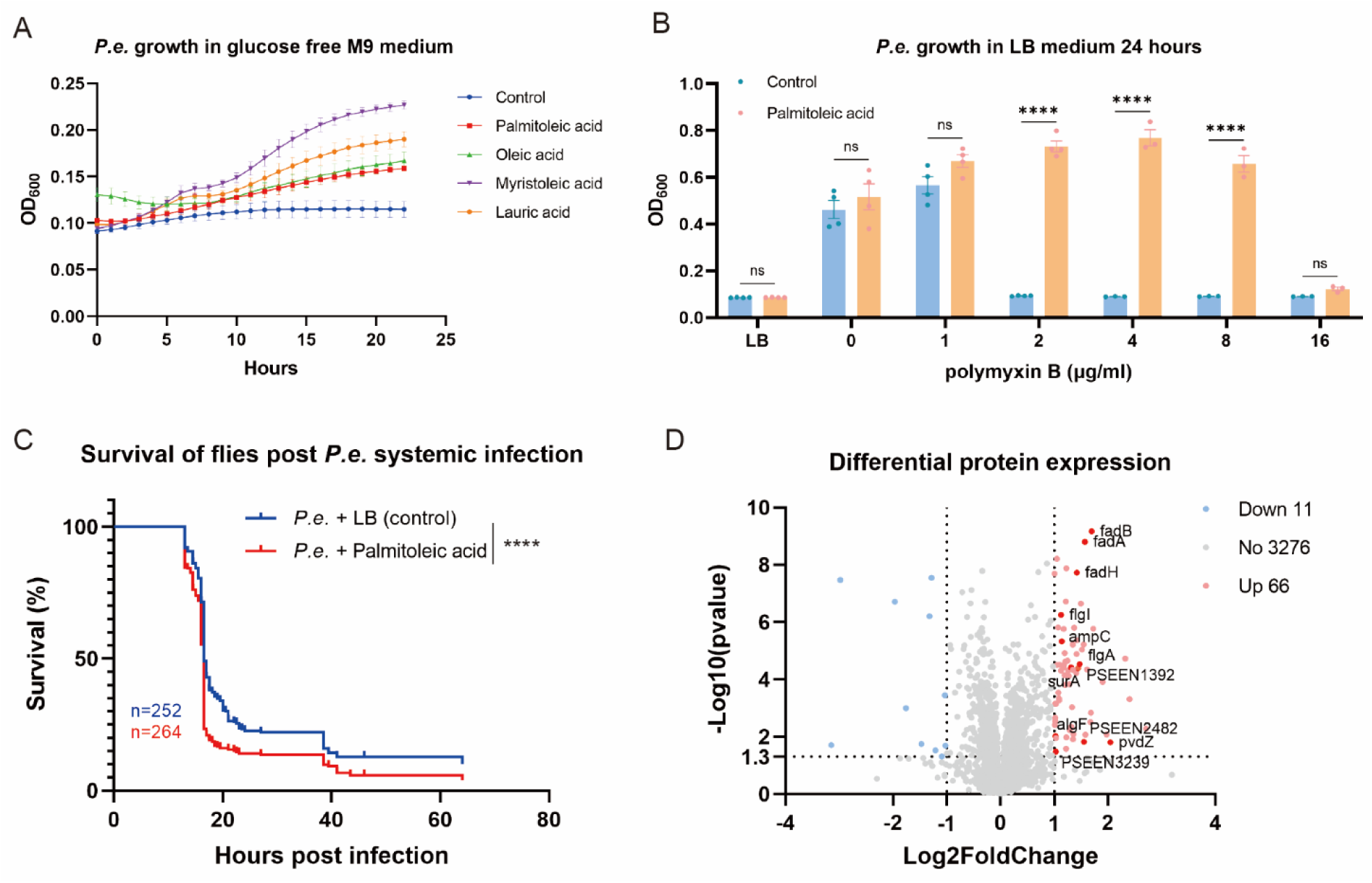
Fatty acids increase *Pe* virulence. (A) Growth curves of *P. entomophila* in glucose-free M9 medium supplemented with different fatty acids (palmitoleic acid 0.5 mM, oleic acid 0.3 mM, myristoleic acid 0.3 mM and lauric acid 0.3 mM) seperately. (B) Growth of *P. entomophila* in LB medium with different concentrations of polymyxin B or with polymyxin B plus 0.5 mM palmitoleic acid. Bars show mean ± SEM. (C) Survival curves of axenic *w^1118^* iso flies systemically infected with *P. entomophila* by pricking or *P. entomophila* pre-incubated with 0.5 mM palmitoleic acid for 3 hours. The accumulated survival graph is shown (*n* = 5). (D) Differential protein expression in *P. entomophila* versus *P. entomophila* pre-incubated with 0.5 mM palmitoleic acid for 3 hours (*n* = 5). Numbers indicate significantly regulated genes (*p* < 0.05 and |Log_2_FoldChange| ≥ 1). Data analyzed with an unpaired t-test (B), stratified Cox hazard ratios (C). ns, not significant; \**p* < 0.05, \*\**p* < 0.01, \*\*\**p* < 0.001, \*\*\*\**p* < 0.0001. For detailed sample sizes and statistical analyses, see Table S1.

To investigate how exposure to palmitoleic acid shapes *Pe* physiology leading to increased virulence, we performed proteomic analysis of *Pe* pre-incubated with palmitoleic acid for 3h in comparison with LB-treated bacteria. We detected 66 proteins with increased and 11 proteins with decreased abundance (log2FC ≥ 1.0) in *Pe* samples treated with palmitoleic acid (Figure 5D, Table S4). As anticipated, fatty acid degradation (Fad) proteins FadA, FadB, FadH showed increased abundance in FA-treated sample. Among proteins with increased abundance, we detected a number of virulence-linked factors, including alginate biosynthesis protein AlgF ^54^; periplasmic chaperone SurA, which modulates virulence and antibiotic resistance ^55^; Glutathione peroxidase PSEEN1392 that protects the bacteria against oxidative stress ^56^; high-affinity zinc uptake system protein ZnuA (PSEEN3239) ^57^; flagellar proteins FlgA and Flgl ^58^. Several proteins involved in siderophore synthesis and uptake were also more abundant in *Pe* treated with palmitoleic acid. Hence, palmitoleic acid might enhance the ability of *Pe* to acquire iron, given the known role of siderophores as high-affinity iron-chelating compounds in iron acquisition ^59,60^. These proteins include one of the pyoverdine synthesis proteins, PvdZ, and two putative TonB-dependent siderophore receptors, PSEEN3587 & PSEEN2482. Beta-lactamase AmpC, conferring resistance to β-lactam antibiotics ^61^, was also more abundant in *Pe* pre-incubated with palmitoleic acid.

Overall, our proteomic analysis demonstrated that palmitoleic acid increases the abundance of proteins involved in virulence and antimicrobial resistance in *Pe*, further supporting the virulence-boosting activity of FAs.

### The ability to utilize FAs is necessary for *Pe* virulence

Considering the virulence-promoting activity of FAs, we hypothesized that blocking the ability of *Pe* to utilize FAs should reduce virulence. Testing this hypothesis requires a mutant that can’t use FAs. To isolate such a mutant, we created a random transposon mutant library, using EZ-Tn5™ <KAN-2>Tnp Transposome™ Kit. We screened part of this library for *Pe* mutants with no/reduced growth in M9 media supplemented with palmitoleic acid as a sole carbon source. In parallel, the growth of all mutants was tested in LB to exclude mutants with generally impaired fitness (Figure 6A). Although our primary screen identified several candidates (Figure 6B), retesting validated 4 mutants that showed comparable growth to wild-type *Pe* in LB (Figure 6C) but impaired growth in M9 medium with palmitoleic acid (Figure 6D). We identified transposon insertion sites in the four mutants and found that the following genes were hit (Figure 6E): *spoT* (guanosine-3’,5’-bis(diphosphate) 3’-pyrophosphohydrolase); *ndk* (nucleoside diphosphate kinase); *PSEEN_RS06965* (putative O-antigen biosynthesis protein); *fadD1* (long-chain-fatty-acid-CoA ligase). Among these genes, *FadD* has been shown previously to be required for bacterial ability to degrade FAs ^62,63^. *P. aeruginosa FadD* mutants have decreased growth on FAs and display impaired virulence in infection models ^62,63^. *spoT* also has an annotated role in FA metabolism as a sensor of fatty acid starvation ^64^. We tested the virulence of the isolated mutants and found that they were unable to kill flies (Figure 6F), suggesting that *Pe*’s ability to utilize FAs is necessary for virulence.

**Figure 6.**
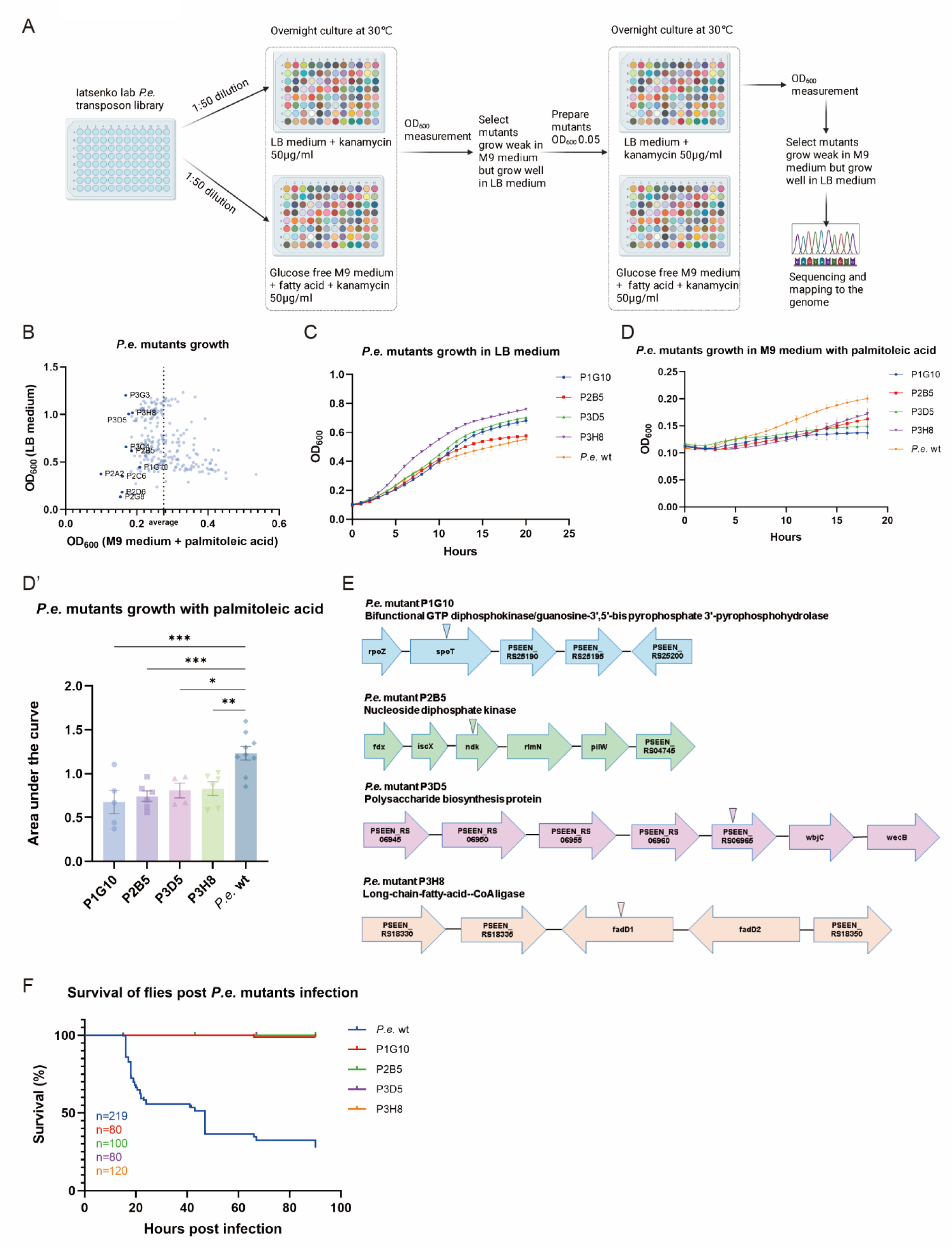
The ability to utilize FAs is necessary for *Pe* virulence. (A) Scheme showing the selection and identification of *P. entomophila* mutants that are deficient in using fatty acids as a carbon source from the transposon library. (B) Scatter plot showing the growth ability of *P. entomophila* mutants in LB medium and M9 glucose-free medium supplemented with palmitoleic acid from the lab transposon library. (C) Growth curves of the wild-type *P. entomophila* strain and selected mutants in LB medium. (D and D’) Growth curves and quantified area under the curve of *P. entomophila* wild-type and selected mutants in M9 glucose-free medium supplemented with palmitoleic acid. Bars show mean ± SEM. (E) Schematic illustration of the genes and their surrounding genomic regions involved in fatty acid utilization. (F) Survival curves of axenic *w^1118^* iso flies infected with wild-type *P. entomophila* and selected mutants. Accumulated survival graph is shown (*n* = 2-3). Data analyzed with one-way ANOVA followed by a Sidak multiple comparisons test (D’). ns, not significant; \**p* < 0.05, \*\**p* < 0.01, \*\*\**p* < 0.001, \*\*\*\**p* < 0.0001. For detailed sample sizes and statistical analyses, see Table S1.

## Discussion

We investigated the role of microbiota in *Drosophila* susceptibility to a natural insect pathogen *Pe* and found that a specific gut commensal, *Lp*, facilitates infection by modulating the host physiology rather than via direct interactions with the pathogen. Our results support the following model (Figure 7). Upon gut colonization, *Lp* enhances the production of several fatty acids, including palmitoleic acid. These FAs support pathogen growth, increase *Pe* virulence and resistance to host antimicrobial effectors, facilitating pathogen persistence in the gut. Persisting pathogens consequently over-stimulate the immune system, leading to over-production of several AMPs. Among these AMPs, Metchnikowin and Attacin D are particularly deleterious and contribute to host mortality during *Pe* infection. This indirect, host-mediated facilitation of infection by microbiota is a sophisticated mechanism that adds a new dimension to our understanding of host-microbe interactions.

**Figure 7.**
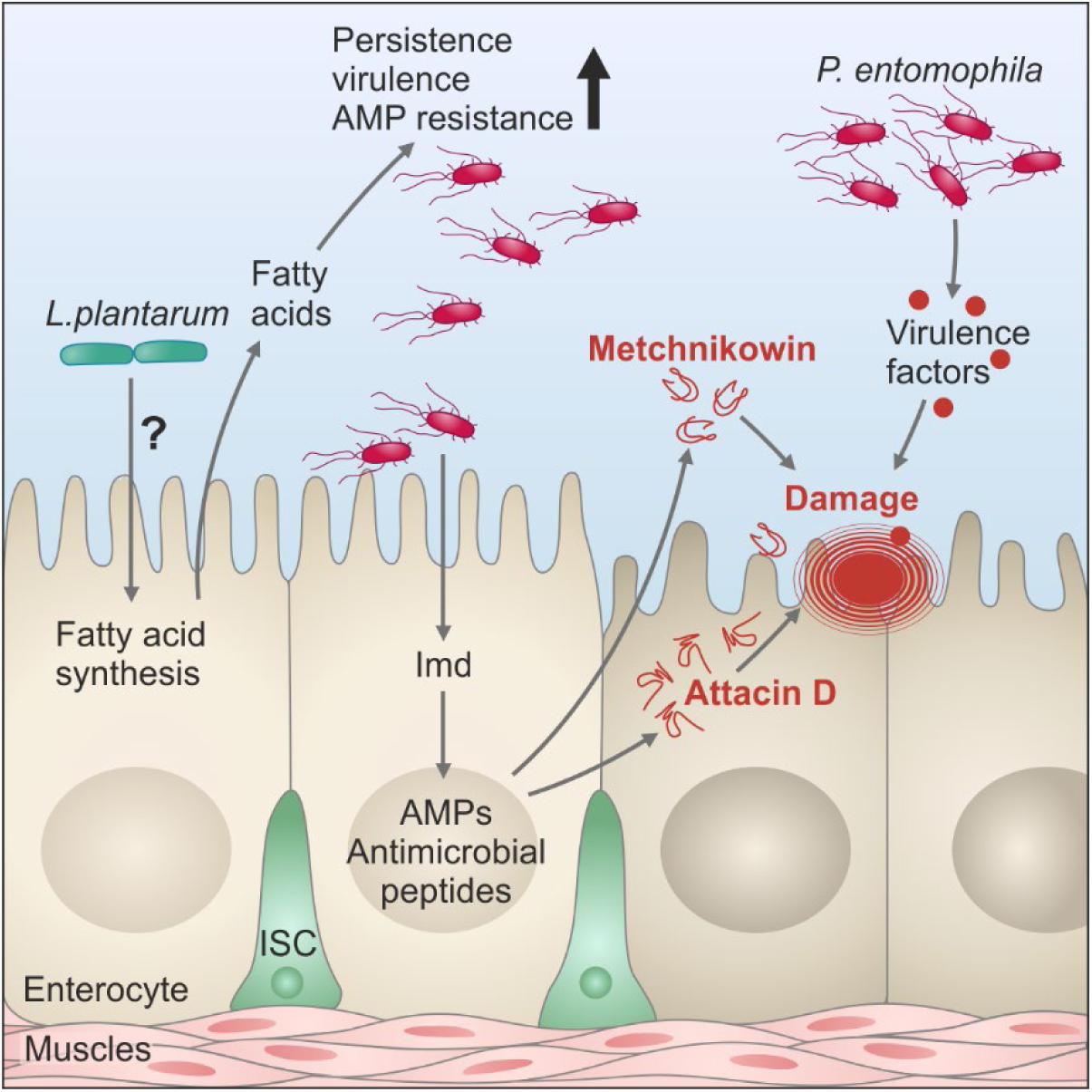
Graphical summary of the mechanism underlying *L. plantarum*-mediated infection facilitation. See text for details.

Our study contributes to the growing body of evidence supporting microbiota-mediated infection facilitation ^5,6^. For instance, the presence of a gut microbiota in flies has been shown to heighten the virulence of *Pseudomonas aeruginosa*, leading to greater mortality among colonized flies compared to germ-free counterparts. Mechanistically, *P. aeruginosa* detects peptidoglycans released from Gram-positive commensals and upregulates its virulence factors in response ^43^. Similarly, the microbiota has been found to exacerbate the pathogenesis of *Vibrio cholerae* in *Drosophila* models ^45^. Importantly, Akyaw et al. described that the mortality of flies following exposure to heat-killed *Pe* is exacerbated in the presence of the microbiota ^65^. Although they used heat-killed pathogen, they found, consistent with our work, that microbiota needs to be present prior to infection. Systemic immune overreaction in microbiota-colonized flies was proposed as a potential mechanism. Our findings, however, contrast with Barron et al., who found the opposite phenotype – host protection against *Pe* infection by *Lp*-mediated acidification ^42^. We believe that differences in the preparation of gnotobiotic flies likely explain the discrepancy in phenotypes. In contrast to Barron et al., who maintained gnotobiotic flies for at least two generations prior to infection, our flies were colonized with *Lp* for only 24 h, which may be insufficient to acidify the gut and confer protection. We can’t exclude that different fly stocks and bacterial strains that we used also contributed to the distinct infection outcomes. Differences between microbial strains are frequently overlooked, yet they may account for some of the inconsistencies observed in the field ^66^.

One of the unexpected findings of our work is increased resistance of *ΔAMP* mutant flies to intestinal infection with *Pe*. AMPs are important immune effectors in flies as illustrated by the remarkable susceptibility of AMP-deficient flies to different pathogens ^50,51,67^. However, so far *ΔAMP* mutant flies have not been tested with intestinal infections. It is commonly assumed that AMPs also contribute to the intestinal defence, given the high susceptibility of Imd pathway mutants that lack inducible AMP expression to intestinal infections ^15^. Our results challenge this assumption. Imd pathway mutant *Relish* consistent with previous studies was more susceptible to *Pe* oral infection ^33^, however, *ΔAMP* mutant flies were more resistant. Hence, susceptibility of Imd pathway mutants to *Pe* is not due to deficiency in AMPs production. What else does Imd pathway control besides AMPs that is necessary to fight *Pe* infection? This will be an important question to address by future studies.

By investigating single AMP mutant flies, we found that most individual AMPs had no significant effect on *Pe* infection. Strikingly, the *Metchnikowin* mutant and RNAi line exhibited remarkable resistance to *Pe* infection, whereas overexpression exacerbated susceptibility. *Attacin D* deficiency led to a similar increased survival of infected flies. AttD and Mtk are likely to cause collateral damage during the over-active immune response. Indeed, accumulating evidence suggests that both AttD and Mtk can cause such damage. For instance, AttD is required for the immune-induced damage of the Malpighian tubules ^68^. Also, AttD deficiency suppressed the defects associated with neurological decline and neurodegeneration in flies with chronic activation of the Imd pathway ^69^. A similar role in neurodegeneration was reported for Mtk. *Mtk* mutant flies show less severe behavioral deficits and increased survival after traumatic brain injury, which increases AMP expression ^70^. Overactivation of *Mtk* was shown to exacerbate poly(GR)-induced neurotoxicity in *Drosophila*, while knockdown of *Mtk* in neurons partially suppresses degeneration ^71^. Hence, under pathological conditions of chronic immune response and tissue damage, some AMPs, like AttD and Mtk can target the host tissues ^72,73^. AttD and Mtk have distinct features that differentiate them from the other AMPs and might explain their contribution to collateral damage. AttD is the only non-secreted AMP in *Drosophila*. It is possible that excessive accumulation of AttD within cells during the immune response can damage organelles. For example, membranes of mitochondria, which are structurally similar to bacterial membranes can potentially be targeted ^74^. Mtk belongs to proline-rich peptides, which operate in a non-lytic manner by blocking the intracellular target, such as succinate-coenzyme Q reductase of fungi or DnaK of bacteria ^75,76^. It is possible that Mtk can’t access these targets in intact cells, however *Pe*-inflicted damage, particularly toxin-mediated pore-formation, can allow Mtk to enter epithelial cells and further exacerbate the damage. Additionally, damaged cells might expose cellular patterns sensitizing these cells to AMPs. For example, Phosphatidylserine (PS) is a negatively charged phospholipid, normally restricted to the inner leaflet of the cell membrane. However, PS can be exposed on the outer leaflet in apoptotic or damaged cells ^77^, hence allowing cationic AMPs to target these cells, as exemplified by specific targeting of tumor cells by Defensin ^78^ or tracheal cells by multiple AMPs ^79^. Further studies are needed to elucidate the exact mechanism of how Mtk and AttD contribute to immunopathology in *Pe* infected flies.

Another important unresolved question is how *Lp* increases FAs synthesis. One possibility is that *Lp* activates insulin/IGF or TOR pathways in the gut or fat body, as was reported in larvae ^80^, promoting anabolic metabolism, including increased fatty acid synthesis. In support of this hypothesis, the expression of *Drosophila insulin-like peptide 3 (Dilp3)* with the known role in promoting IIS activity ^81^ and fat storage ^82^, is induced by *Lp* colonization. Another possibility can be that *Lp* produces metabolites that act as building blocks for FAs or as signaling molecules, triggering increased expression of lipogenic enzymes in the host. For example, acetyl-CoA, a crucial precursor of FAs, can be generated from *Lp*-derived lactate ^83^. Elucidating the mechanisms by which *Lp* induces FAs synthesis and how these fatty acids affect *Lp* itself, as was reported for other commensals ^84^, remain important areas for investigation. Given our findings that *Lp* abundance increases in infected flies ^85–87^, fatty acids could contribute to this effect by enhancing *Lp* resistance to AMPs.

Overall, our work uncovers an exemplary case of microbiota-mediated changes in host physiology that predispose the host to enteric infection. The protective or detrimental role of the microbiome is often context- and pathogen-dependent, necessitating more direct empirical investigations to distinguish the conditions under which the microbiota might facilitate infection. The *Drosophila* model of host-microbiome-pathogen interactions offers an accessible genetic system for understanding the mechanistic links between microbiome-modulated host physiology and infection outcome.

## Materials and Methods

### Drosophila husbandry

*Drosophila* stocks were raised in a light:dark cycle (12 h:12 h) at 25°C, on a standard cornmeal/agar medium (6.2 g agar, 58.8 g cornmeal, 58.8 g inactivated dried yeast, 26.5 mL of a 10% solution of methyl-paraben in 85% ethanol, 60 mL fruit juice, 4.8 ml 99% propionic acid for 1 L). Flies were transferred to fresh vials weekly to maintain them. Adult flies were allowed to mate for 2-3 days after eclosion. Then, flies were sorted in vials under CO_2_ with 20 flies per vial before the experiment. Axenic flies were generated by egg bleaching as previously described ^24,86^. The food for the axenic flies contained ampicillin (50 µg/mL), kanamycin (50 µg /mL), tetracycline (10 µg/mL), and erythromycin (10 µg /mL). Fresh food was prepared weekly to avoid desiccation.

### *Drosophila* strains and crosses

*Drosophila* strains used in this study are listed in Table S5. To prepare the crosses, 5 males carrying UAS transgenes were placed together with 10 virgin females carrying the GAL4 driver. The flies were transferred to fresh vials every 2-3 days at 25°C. After eclosion, all emerged adults were kept for 2 days to allow mating. Then, the flies were transferred to 29°C for at least 3 days before the experiments. Female flies were used for all experiments.

### Isolation of bacteria from conventional flies

Conventional wild-type *Drosophila melanogaster* flies were maintained on standard cornmeal–yeast–agar medium at 25 °C. Adult flies (around 10 days old) were anesthetized with CO₂ and surface-sterilized by washing in 70% ethanol for 1 min followed by two rinses in sterile phosphate-buffered saline (PBS). Groups of 5 flies were homogenized in 500 µL sterile PBS using sterile plastic pestles. The homogenates were plated onto LB, Mannitol, and MRS agar plates. The plates were then incubated aerobically at 30 °C for 24–72 hours, after which individual colonies were selected based on morphology and re-streaked to obtain pure cultures. Bacterial isolates were initially identified using MALDI-TOF mass spectrometry (VITEK MS, bioMérieux). Briefly, individual colonies from freshly streaked plates were transferred onto a MALDI target plate and overlaid with matrix solution according to the manufacturer’s protocol. Spectra were acquired and compared against the reference database for species-level identification.

To confirm the taxonomic assignment, genomic DNA was extracted from the bacterial cultures, and the 16S rRNA gene sequencing was performed using universal bacterial primers (8F: 5’-AGAGTTTGATCCTGGCTCAG-3’; 1492R: 5’-GGTTACCTTGTTACGACTT-3’). The PCR products were purified and subjected to Sanger sequencing. The resulting sequences were analyzed using BLAST against the NCBI database to determine taxonomic identity. Using this approach, the isolates were identified as common fly-associated bacteria, including *Lactiplantibacillus plantarum* and *Levilactobacillus brevis*. The bacterial strains used in this study are listed in Table S5.

### Oral infection and survival assay

Unless otherwise indicated, the previously established protocol (Figure 1A) was used for the oral infection of flies^88,89^. The *Drosophila* natural pathogen *P. entomophila* L48 strain was obtained from Bruno Lemaitre’s laboratory. The *P. entomophila* strains used in this study were grown directly from frozen 50% glycerol stocks. Briefly, 50 μL of the stock was incubated in 20 mL of LB medium in a 50 mL flask overnight (∼16 h) at 30°C with shaking at 175 rpm. The following day, the overnight culture was diluted 1:15 in 150 mL of LB medium and incubated in flasks under the same conditions for a minimum of an additional 24 hours. Before the infection experiment, the cultures were centrifuged at 3,500 rpm and 15°C for 15 minutes. Their concentration was adjusted to the optical density (OD_600_) of 200 with PBS. The bacterial suspension was then mixed 1:1 with a 5% sucrose solution before the infection experiment. Then, 150 μL of the mixture was pipetted into vials containing standard food and filter papers (referred to as infection vials). Prior to transferring mated female *w^1118^* iso axenic flies to each infection vial, the flies were kept in empty vials for 2 hours at 29°C (dry starvation) (Fig. 1A). The flies were kept in the infection vials for 24 hours before being transferred to conventional fly food vials. All infections were performed at 29°C. The number of dead flies was counted continually at the different time points. All experiments were performed with multiple biological repeats, each repeat with at least two vials containing approximately 20 flies.

### *Pe* systemic infection

The *Pe* culture and *Pe* pre-incubated with individual fatty acids (palmitoleic acid 0.5mM, myristoleic acid 0.3mM or lauric acid 0.3mM, all fatty acids were from Sigma-Aldrich) for 3 hours were pelleted by centrifugation to remove the media. Then, they were diluted to the desired optical density (OD_600_ = 1) with sterile PBS. To infect axenic *w^1118^* iso flies, a 0.15 mm minutien pin (Fine Science Tools) mounted on a metal holder was dipped into the diluted bacterial solution and poked into the thorax of a CO_2_ anesthetized fly. Infected flies were maintained in vials (20 flies per vial) with food at 29 °C. Surviving flies were counted at regular intervals. One to three vials were infected per experiment, with at least two experiments performed.

### Commensal bacterial mono-inoculation assay

Commensal bacteria were cultured in a suitable medium for one or two days before harvesting for mono-inoculation in flies. *L. plantarum WJL*, *L. plantarum YY*, and *Levilactobacillus brevis* were cultured in MRS medium at 37°C with shaking at 175 rpm. *Sphingomonas leidyi*, *Sphingomonas sp.,* and *Variovax sp.* were cultured in mannitol medium at 30°C with shaking at 175 rpm. *Acetobacter malorum* and *Acetobacter pomorum* were cultured in MRS medium at 30°C with shaking at 175 rpm. The cultures were then centrifuged at 3,500 rpm at15°C for 15 minutes. Their concentration was estimated to the optical density (OD_600_) at 50 using PBS. The bacterial suspension was then mixed at a ratio of 1:1 with a 5% sucrose solution. Next, 150 μL of this mixture was pipetted into vials containing standard food and filter papers (Fig. 1A). Flies were starved in empty vials for 2 hours at 29°C (dry starvation) before flies were transferred to the vials containing commensal bacteria. The colonization time is 24 hours unless otherwise specified.

### Bacterial loads measurement

To measure bacterial load in flies, 5 female flies from each vial were collected at 3, 6, and 9 hours post *Pe* infection. The samples were first surface-sterilized in 70% ethanol for 20 seconds, with three washes and one in sterile PBS for 30 seconds. Then, the samples were homogenized in 500 μL of sterile PBS for 30 seconds at 6000 rpm using a Precellys 24 instrument (Bertin Technologies, France). Serial 10-fold dilutions ranging from 10⁻^1^ to 10⁻^6^ were made, and 10 μL of each dilution with 3 replicates were plated on LB plates. After an overnight incubation (proxy 18 h) at 30°C, the number of colonies were counted. Flies fed with sucrose were processed in parallel with infected flies as a control. Since the control plates were negative for bacterial colonies, it was assumed that all colonies that grew on plates from infected flies were *P.e*. To measure bacterial loads in an *in vitro* culture, a starting OD_600_ of 0.05 was prepared for *Pe* and *Lp YY* in BHI medium. After 24 hours, serial 10-fold dilutions ranging from 10⁻^1^ to 10⁻^10^ were made, and 10 μL of each dilution with 3 replicates were plated on LB plates. The following procedure is the same as the bacteria loads measurement in flies.

### Food intake quantification

The amount of ingested food was quantified by including blue dye (Food Blue No. 1, TCI) in the infection mixture modified from a previous method ^89^. *Pe* was mixed at a 1:1 ratio with a 5% sucrose solution containing 1% blue dye. After 1 and 2 hours, feeding was interrupted and five flies were transferred to tubes containing glass beads and 500 μL of 1x PBS with 0.1% Triton X-100 (PBST). The samples were then homogenized using a Precellys homogenizer (three cycles of 30 seconds at 7,200 rpm) and then centrifuged twice (2 mins, maximum speed). Then, 300 μL of the supernatant was then transferred to a cuvette containing 700 μL of water. Blue dye consumption was quantified by measuring absorbance at 630 nm using an Ultrospec 2100 pro UV/Vis Spectrometer (Amersham Biosciences).

### Measuring defecation in adult flies

Ten female flies were kept in each vial containing an infection mix (*Pe* and 2.5% sucrose) or a control (2.5% sucrose) and blue dye as described above (see: “Food intake quantification”). The flies were kept on the blue suspension for 2 hours and then transferred to normal food vials for an additional 2 hours. The flies were transferred to normal food vials again for 16 additional hours. Defecation dots from the first 4 hours and a total of 20 hours were quantified. The defecation rate was measured by counting the “defecation dots” left dried on the inner walls of the vials and normalized the count by per number of flies in each vial.

### Hemolymph bacteria detection

Axenic *w^1118^* iso flies were either colonized with *Lactiplantibacillus* or orally infected with *Pe* under the following conditions (1) 2.5% sucrose 24 hours, followed by sucrose for 13 hours; (2) *Lp YY* colonized for 24 hours, followed by sucrose for 13 hours; (3) 2.5% sucrose 24 hours, followed by *Pe* infection for 13 hours; (4) *Lp YY* colonized for 24 hours, followed by *Pe* infection for 13 hours; (5) positive control: systemic infection of *Pe* OD_600_ = 1. Colonization and oral infection conditions were the same as previously mentioned (see “Oral infection and survival assay”).

The hemolymph extraction method from flies was modified from the previous method^90^, Briefly, the glass capillary was first prepared. The capillary puller heater (Narishige PC-100 Vertical puller, Washington D.C., USA) was set to 55% of maximum setting. The capillary tube was pulled into a sharp point with an approximate diameter of 10 µm. The flies were then anesthetized with CO2 gas for 5 s, and the glass capillary was lightly placed onto the thorax to absorb the hemolymph. Hemolymph from around 30 flies was pooled into 250 µL of sterile PBS, after which 100 µL was plated on MRS and LB plates separately. The bacterial colonies that grew on the MRS and LB plates were photographed by using the Scan 300 Automatic colony counter (Interscience, France).

### Transcriptomic analysis

We characterized the transcriptional changes in the fly gut under the following treatment conditions: (1) 2.5% sucrose 24 hours, followed by sucrose for 16 hours; (2) *Lp YY* colonization 24 hours, followed by sucrose for 16 hours; (3) 2.5% sucrose 24 hours, followed by *Pe* for 16 hours; (4) *Lp YY* colonization 24 hours, followed by *Pe* for 16 hours. *Lp YY* was prepared as previously mentioned. Total RNA was extracted from 20 guts per sample using TRIzol reagent (Invitrogen). The fly guts were dissected on ice and homogenized in 500 µL of Trizol plus 150 µL of chloroform for 30 seconds at 7,200 rpm with beads. Then, the typical RNA extraction procedure was followed. The quantity and quality of RNA were measured using a NanoDrop 2000 (Thermo Scientific). For each condition, RNA was prepared from three biological replicates on different days. RNA-seq was carried out by Novogene (Cambridge, UK) using standard Illumina protocols. Briefly, the quantity, integrity, and purity of the RNA were determined using the Agilent 5400 Fragment Analyzer System (Agilent Technologies), and the mRNA was purified from the total RNA using poly-T oligonucleotide-attached magnetic beads.

After fragmentation, the first-strand cDNA was synthesized using random hexamer primers. Then, the second-strand cDNA was synthesized using dUTP, instead of dTTP. After end repair, A-tailing, adapter ligation, size selection, USER enzyme digestion, amplification, and purification, the directional library was ready. The libraries were quantified with Qubit and checked with bioanalyser for size distribution. The quantified libraries were pooled and sequenced on the Illumina NovaSeq 6000 platform (2 × 150 bp), generating approximately about 6 Gb of raw data per sample.

The differential expression analysis was performed using DESeq2^91,92^. Differential expression analysis of two conditions/groups (3 biological replicates per condition) was performed using the DESeq2R package (version 1.20.0). DESeq2 provides statistical routines for determining differential expression in digital gene expression data using a model based on the negative binomial distribution. The resulting p-values were adjusted using the Benjamini and Hochberg’s approach for controlling the false discovery rate. Genes with an adjusted P-value ≤ 0.05, as determined by DESeq2, were considered differentially expressed. Volcano plots were made using a cut-off of log2FC ≥1 to visualize genes that were upregulated or downregulated compared to the appropriate groups. Gene Ontology^93^ (GO) enrichment analysis of differentially expressed genes was implemented by the Cluster Profiler R package, which corrected for gene length bias. GO terms with a corrected P-value ≤ 0.05 were considered to be significantly enriched by the differentially expressed genes. All raw RNA sequencing data files are available from the SRA database (accession number PRJNA1435853) and can be accessed at https://www.ncbi.nlm.nih.gov/sra/PRJNA1435853.

### Measurement of free fatty acids in fly guts

Free fatty acids were measured in the fly guts under the following treatment conditions: (1) 2.5% sucrose for 24 hours; (2) *Lp YY* colonization for 24 hours, or (3) *Acetobacter* colonization for 24 hours. The *Lp YY* and *Acetobacter* were prepared as previously mentioned. The weight of individual Eppendorf tubes was measured before sample collection. Twenty guts were dissected on ice and pooled into 500 µL of cold PBS in a tube. The tubes were centrifuged at 12,000 rpm for 1 min at 4 °C, then the liquid was completely removed. The tubes were centrifuged a second time, if necessary, to completely remove the liquid. The weight of each tube was measured again. The weight difference indicates the weight of the collected fly guts, which was used to normalize the level of free fatty acids. Free fatty acids were extracted from the gut samples according to the instructions provided in the Free Fatty Acid Quantitation Kit (catalog number MAK044, Sigma-Aldrich). In brief, the fly guts were homogenized in a solution of 200 µL of a 1% (w/v) Triton X-100 in chloroform. The samples were centrifuged at 13,000 g for 10 minutes to remove insoluble material. The organic phases (lower phase) were collected and air-dried at 50 °C to remove the chloroform. They were vacuum dried for 30 minutes to remove any remaining chloroform. The dried lipids were dissolved in 200 µL of Fatty Acid Assay Buffer by extensive vortexing for 5 minutes. Palmitic acid standards from the kit were prepared for colorimetric detection. The samples were brought to a final volume of 50 µL with the fatty acid assay buffer. After performing the assay reaction with the master reaction mix, colorimetric assay was used to measure the absorbance at 570 nm (A570). The concentration of fatty acids was calculated based on the standards and normalized based on the weight of the gut (mg). The relative levels of fatty acids in the commensal bacteria-colonized fly guts, compared to the control group, were plotted.

### Protease inhibition in flies

AEBSF protease Inhibitors (0.25, 0.5, and 1mM) were mixed with *Lp YY* to colonize the axenic flies. Sucrose (2.5%) containing the same concentration of AEBSF was used as the controls. The bacteria colonization method was introduced in the previous section.

### Lipidomic analysis

#### Sample Preparation

Lipidomic analysis was performed by Lipotype (Dresden, Germany) on whole flies with the following treatment conditions: (1) 2.5% sucrose for 24 hours, followed by 4 hours of 2.5% sucrose; (2) *Lp YY* colonization for 24 hours, followed by 4 hours of 2.5% sucrose. *Lp YY* was prepared as previously mentioned. Approximately 20 mg of whole flies were weighed precisely, then homogenized in 500 µL of isopropanol using TissueLyzer III (Qiagen) for 15 minutes at 30 Hz. The homogenate was treated in an ultrasonic bath for 10 minutes and then centrifuged for 5 min at 11,000 rpm. A 50 µL aliquot was spiked with a mixture of deuterated internal standards, consisting of: (1) C6:0-d11, (2) C12:0-d23, (3) C18:0-d35, (4) C18:2-d4, (5) C20:4-d11, (6) C20:5-d5, (7) C22:0-d43, (8) C22:6-d5, and (9)C24:0-d4 with 10 ng each (Cayman Chemical, Ann Arbor, USA), in a 450 µL Methanol solution in an autosampler glass vial.

#### LC/ESI-MS/MS

The clear solutions were analyzed using an Agilent 1290 HPLC system with a binary pump, a multisampler and a column thermostat with a Kinetex C-18 column (2.1 x 150 mm, 2.7 µm). A gradient solvent system of aqueous acetic acid (0.05 %) and acetonitrile was used. The flow rate was set at 0.4 mL/min and the injection volume was 1 µL. The HPLC system was coupled with an Agilent 6470 Triplequad mass spectrometer (Agilent Technologies, Santa Clara, USA) with an electrospray ionization source. Analysis was performed with Multiple Reaction Monitoring in negative mode, with at least two mass transitions for each compound. All fatty acids were individually calibrated using a 37-component Supelco FAME-Mix (Merck, Darmstadt, Germany) in relation to deuterated internal standards.

### Bacterial growth dynamic in M9 medium supplemented with fatty acids

The *Pe* growth dynamics was tested in glucose-free M9 medium supplemented with individual fatty acids (palmitoleic acid 0.5 mM, oleic acid 0.3 mM, myristoleic acid 0.3 mM and lauric acid 0.3 mM) and using the same amount of solvent (100% ethanol) as a control. The *Pe* was cultured and pelleted as previously mentioned, and then diluted to an OD_600_ = 0.05 in M9 medium with the individual supplements. To prepare 100 mL of M9 medium, 20 mL of M9 salts (5X), 2 mL of glucose (20%; Sigma-Aldrich, filter-sterilized and stored at 4°C), 200 μL of MgSO₄ (1 M; Fisher Scientific, autoclaved and stored at room temperature), 10 μL of CaCl₂ (1 M; Fisher Scientific, autoclaved and stored at room temperature), and 78 mL of H₂O were used. Glucose was excluded when preparing the glucose-free M9 medium.

### Minimum inhibitory concentration (MIC) of polymyxin B

Overnight *Pe* cultures were adjusted to an OD600 of 1. Then, 25 μL of the bacterial suspension was added to 0.5 mL of LB or M9 medium (with glucose) containing different concentrations of polymyxin B (0, 1, 2, 3, 4, 5, 6, 8, or 16 μg/mL) (Fischer Scientific). Fatty acids (final concentrations of palmitoleic acid 0.5 mM, oleic acid 0.3 mM, myristoleic acid 0.3 mM, and lauric acid 0.3 mM) were added and mixed well to test their influence on *Pe* resistance against polymyxin B. The corresponding amount of solvent (100% ethanol) was used as a control. One hundred μL of the suspension was pipetted into 96-well plates. After an overnight incubation, the bacterial growth dynamics (OD_600_) was recorded using an Infinite M Plex Microplate Reader (Tecan, Austria) to determine MIC value.

### Bacterial proteomic analysis

#### Samples preparation for proteomics

Bacteria were cultured overnight and diluted to an O_D600_ of 1. *Pe* was incubated with palmitoleic acid (0.5 mM) or with solvent control in the volume of 5 mL for 3 hours. Bacteria were centrifuged at 3,500 rpm and 4 °C for 10 minutes. After centrifugation, the LB medium was discarded completely. The bacterial pellet was resuspended and washed with 600 μL of extraction buffer without SDS (100 mM Tris-Cl pH 8 + 1X Roche protease inhibitor). The bacteria were collected by centrifugation at 3,500 rpm, 4 °C for 5 min, and the washing buffer was discarded. Then, 500 μL of extraction buffer (100 mM Tris-Cl pH 8, 2% SDS, and 1X Roche protease inhibitor) was added to the pellet, which was then resuspended. The suspension was incubated at 95°C for 2 min, then homogenized in a Precellys 24 Tissue Homogenizer at 6,000 rpm for 30 seconds. The suspension was centrifuged for 5 min at 12,000 rpm and 4°C, and the supernatant was collected in a new Eppendorf tube. The protein concentration was measured using the Pierce BCA Protein Assay Kit (Thermo Fisher) according to the manufacturer’s protocol. Samples were stored at -80°C.

#### Sample analysis by LC-MS

All samples were subjected to the SP3 sample preparation protocol ^94^. Ten µg of a 1:1 mixture of hydrophilic and hydrophobic carboxyl-coated paramagnetic beads (SeraMag, #24152105050250, GE Healthcare) were added for each µg of protein. Protein binding was induced by the addition of acetonitrile to a final concentration of 70% (v/v). Samples were incubated at room temperature for 10 minutes. The tubes were placed on a magnetic rack, and beads were allowed to settle for three minutes. The supernatant was discarded, and beads were rinsed thrice with 80% ethanol. Beads were resuspended in a digestion buffer containing 50 mM triethylammonium bicarbonate (Sigma, #T7408) and Trypsin and lys C (SERVA, #37283.03) in a 1:50 enzyme-to-protein ratio. Protein digestion was carried out for 14 hours at 37°C. Afterward, the peptide supernatant was recovered and acidified with 2% ACN and 0.1% trifluoroacetic acid.

Label-free DIA analyses of peptides were acquired over 120 min by an Orbitrap Exploris 480 (ThermoScientific) coupled to a 3000 RSLC nano UPLC (ThermoScientific) from 750 ng of peptides. Samples were loaded on a PepMap trap cartridge (300 µm i.d. × 5 mm, C18, ThermoScientific) with 2% acetonitrile, 0.1% TFA at a flow rate of 20 µL/min. Peptides were separated over a 25 cm analytical column (PepSep C18, 75 µm I.D., 1.5 µm). Solvent A consists of 0.1% formic acid in water. Elution was carried out at a constant flow rate of 250 nL/min within 120 min. Initially, a two-step linear gradient was applied: 5–30% solvent B (0.1% formic acid in 80% acetonitrile) over 70.5 min, then 30–45% solvent B over 13 min, followed by column washing and equilibration. The column was kept at a constant temperature of 50 °C.

The MS was operated in DIA mode for single-injection quantitative measurements of individual samples with the following settings: 60k MS1 resolution, MS1 scan range 350–1250 m/z, 15k MS2 resolution, MS2 scan range 110–1600 m/z, Normalised AGC target of 1000%, maximum injection time 60 ms, and fixed normalized collision energy of 30. DIA MS2 scans were performed at 12 m/z precursor isolation windows with optimized window placements from 400.4319 to 1204.7975 m/z within the precursor mass range 400-1200 m/z with isolation window overlap 0.2 m/z.

Raw data analysis was performed using Spectronaut (Biognosys AG, Zurich, Switzerland) version Spectronaut 20.3.251119.92449 in direct DIA+ deep mode with reviewed UniProt databases (*Pseudomonas entomophila* (strain L48) - UP000000658, 5,126 entries in UniProtKB). Methionine oxidation and Acetyl (Protein N-term) were set as a variable, and carbamidomethylation on cysteine residues was used as a static modification. The FDR for PSM-, peptide-, and protein-level was set to 0.01. All tolerances were set to dynamic for pulsar searches. The mass spectrometry proteomics data have been deposited to the ProteomeXchange Consortium via the PRIDE ^95^ partner repository with the dataset identifier PXD076700. Further data processing was carried out in R (v4.5.2) and Perseus (v. 2.0.6.0). Normalized abundances were log2-transformed and filtered to retain proteins with at least 4 valid values in at least one group; otherwise, the data were discarded. Missing values were imputed following normal distribution (0.3 (width), shift it down by “1.8” (down shift) standard deviations

### Generation and screening of a random transposon mutant library in *Pe*

The random transposon mutant library in *Pe* was generated using the EZ-Tn5™ <KAN-2>Tnp Transposome™ Kit (Cat. No. TSM99K2) as described previously ^96^. Electrocompetent cells of *Pe* were prepared by collecting the cells when they reached OD 0.5 and washing them 3 times with cold 10% glycerol. 100 μl of electrocompetent cells were mixed with 1 μl of transposome complex and placed in 0.2 mm electroporation cuvette (Biorad). The cells were electroporated using Gene Pulser Xcell System (Biorad) with the following settings 2.5 kv, 200 ohms and 25 μF. Cells were recovered immediately following the pulse with 1 ml of LB media and incubated for 1 h at 37°C before plating on LB agar plates with 50 μg/ml kanamycin. Following overnight incubation at 37°C, individual colonies were picked into the wells of 96 deep-well plates containing 1 ml of LB+kanamycin per well. The plates were covered with air-permeable sealing films and incubated at 37°C overnight. Next day, 750 μl of 50% glycerol were added to each well and the plates were placed at -80°C freezer for storage.

### Screening for *Pe* mutants deficient in metabolizing fatty acid

*Pe* transposon mutants were recovered from -80°C. 150 μL of LB medium containing 50 μg/mL kanamycin was pipetted into each well of a standard 96-well plate. The wells were inoculated with 3 μL transposon mutants using a multichannel pipette. The bacteria were grown overnight (or longer) stationarily at 30°C. 150 μL of glucose-free M9 medium containing 50 μg/mL kanamycin and 0.5 mM palmitoleic acid was pipetted into each well. The wells were inoculated with 3 μL transposon mutants using a multichannel pipette. The plates with either LB medium or M9 medium were incubated at 30°C. Bacterial growth was measured with OD600 after 24 hours. ***Pe*** mutants that grew normally in LB medium but were impaired in growth in M9 medium plus palmitoleic acid were selected. The selected mutants were then validated using the same procedure, but with a controlled bacterial inoculum (OD600 = 0.05). The validated mutants with reduced ability to metabolize fatty acid were stored as glycerol stocks for further experiments.

### Identification of transposon insertion sites

Genomic DNA was extracted from *Pe* mutants of interest using a Monarch Genomic DNA Purification Kit (New England Biolabs, NEB). The DNA was randomly fragmented with NEBNext dsDNA Fragmentase (NEB). The fragmented DNA was end-repaired (made blunt-ended) and 5′-phosphorylated using an End-It DNA End-Repair Kit (Lucigen). Finally, the DNA was self-circularized using T4 DNA Ligase (ThermoFisher) and transformed into TransforMax EC100D pir+ *E. coli* (Lucigen), which expresses the pir gene product (“pi” protein). When grown on kanamycin-containing plates, only cells containing the <R6Kγori/KAN-2> transposon can grow. Plasmids were extracted from the resulting colonies using Monarch Plasmid Miniprep Kit (NEB) and were used to sequence the transposon-flanking DNA with transposon-specific primers supplied with the kit. Blast search of the obtained sequences was performed to determine the identity of the transposon-disrupted genes.

### Quantification and statistical analysis

The statistical parameters and tests are shown in the figure legends and Table S1. Although no formal randomization method was used, we varied the order of sorting flies, infection vials, sampling and well plate location of samples across different biological replicates to reduce potential bias. Additionally, the control and treatment groups were processed in parallel, and the sample sizes were balanced. Blinding was not implemented in this study. Data analyses were performed using GraphPad Prism 9.5 and R (v4.5.2) software. The kinetics of the survival data are shown using cumulative data, and the survival analysis was performed using a stratified Cox proportional hazards model using the Survival R package. Log hazard ratios are presented to estimate the difference in survival considering all the parameters. The Survival package provided 95% confidence intervals, which were approximated by calculating 1.96 times the standard error. Data visualization was performed using the GraphPad Prism 9.5. Statistical significance was determined using an unpaired Student’s t-test or a one-way or two-way ANOVA, followed by post-hoc tests for multiple comparisons (as indicated in the figure legends). Significance was set at p < 0.05, and data are presented as mean ± SEM. Significance: *p < 0.05, **p < 0.01, ***p < 0.001, ****p < 0.0001.

## Acknowledgments

We are grateful to the Vienna Drosophila Resource Center and the Bloomington Drosophila Stock Center (NIH P40OD018537) for fly stocks. We thank Florian Kondrot (MPUSP) for providing technical support for the mass spectrometry experiments, Diane Schad for help with preparation of graphical abstract, and Dr. Dhiren Patel for fruitful discussions. This work was supported by the Max Planck Society. E.C. acknowledges the Deutsche Forschungsgemeinschaft (Leibniz Prize). I.I. also acknowledges the funding from the Deutsche Forschungsgemeinschaft (grants IA 81/2-1 and IA81/3-1) and from the Boehringer Ingelheim Foundation. YY was supported by a fellowship from the Alexander von Humboldt Foundation.

## Author Contributions

Investigation, formal analysis, data curation, validation, methodology, visualization, writing – original draft preparation, and writing – review and editing, Y.Y.; investigation, validation, visualization, formal analysis, and writing – review and editing, K.A.; investigation and resources, D.F.; funding acquisition, resources, writing – review and editing, E.C.; conceptualization, funding acquisition, project administration, supervision, resources, writing – original draft preparation, and writing – review and editing, I.I.

## Declaration of Interests

The authors declare no competing interest.

---

**Figure S1.**
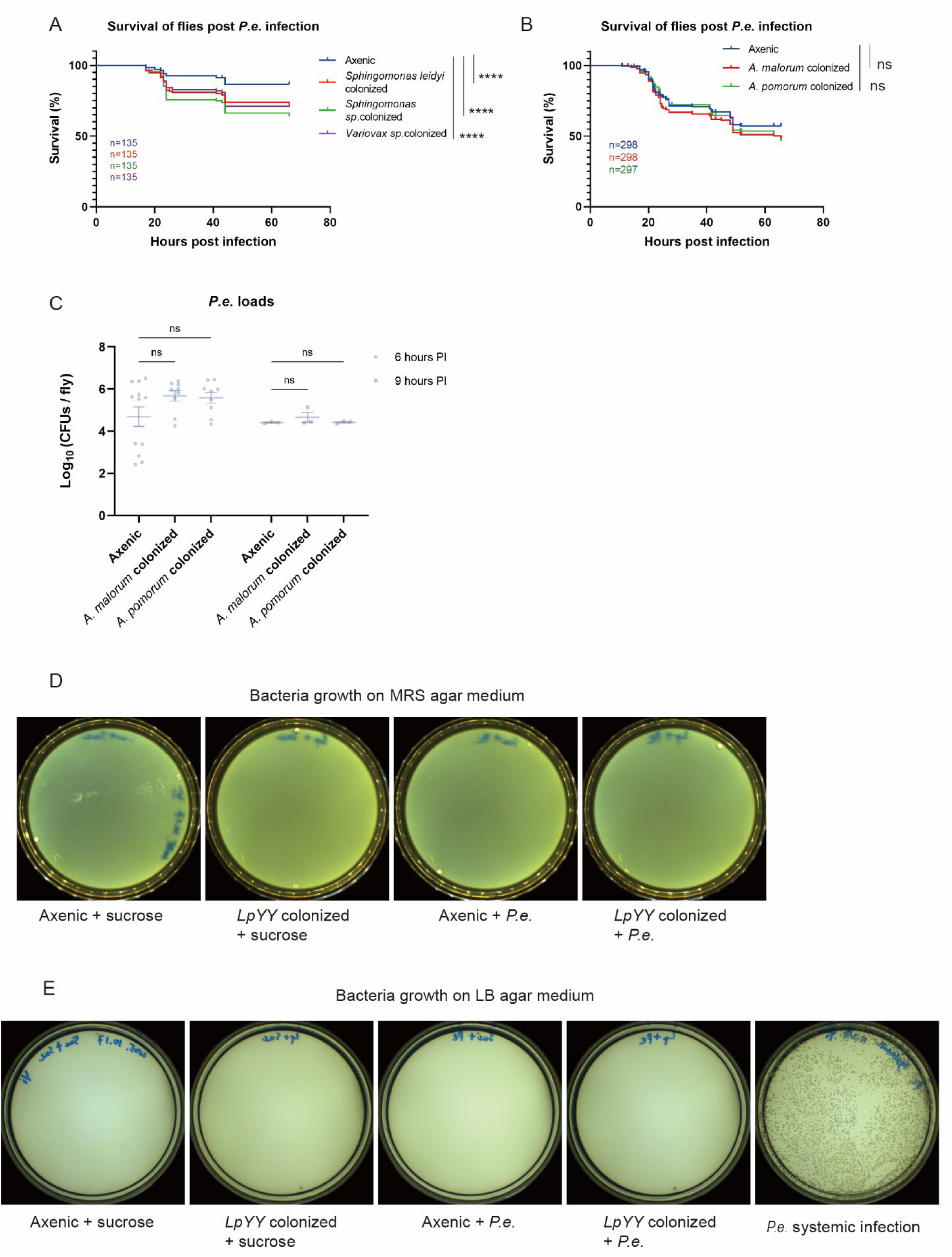
Effect of gut microbiota on fly susceptibility to *Pe* infection and pathogen translocation into hemolymph. (A and B) Survival curves of axenic *w^1118^* iso flies colonized with gut microbiota upon *P. entomophila* infection. The following microbiota species were used: *Sphingomonas leidyi* (A)*, Sphingomonas sp.* (A), *Variovax sp.* (A), *A. malorum* (B), and *A. pomorum* (B). The accumulated survival graph is shown (A, *n* = 3; B, *n* = 7). (C) Pathogen loads (CFUs) at 6 and 9 hours post-exposure to *P. entomophila* in axenic and *Acetobacter* colonized *w^1118^* iso flies. Each dot indicates a pool of five flies taken from different infection vials (*n* ≥ 3, pooled from at least 1 experiments). Bars show mean ± SEM. (D and E) Bacteria detection in hemolymph collected from around 30 axenic *w^1118^* iso flies on MRS (D) and LB (E) agar medium. Hemolymph samples were detected from: (1) unchallenged axenic flies with a 12-hour sucrose treatment (axenic + sucrose) as a negative control; (2) *Lactiplantibacillus*-colonized flies with a 12-hour sucrose treatment (*LpYY*-colonized + sucrose); (3) axenic flies with a 12-hour *P. entomophila* infection (axenic + *P.e.*); (4) *Lactiplantibacillus*-colonized flies with a 12-hour *P. entomophila* infection (*LpYY*-colonized + *P.e.*); (5) axenic flies upon *P. entomophila* systemic infection via pricking as a positive control (*P.e.* systemic infection). Data analyzed with stratified Cox hazard ratios by R (A and B) and two-way ANOVA followed by Sidak’s multiple comparisons test (C). ns, not significant; \**p* < 0.05, \*\**p* < 0.01, \*\*\**p* < 0.001, \*\*\*\**p* < 0.0001. For detailed sample sizes and statistical analyses, see Table S1.

**Figure S2.**
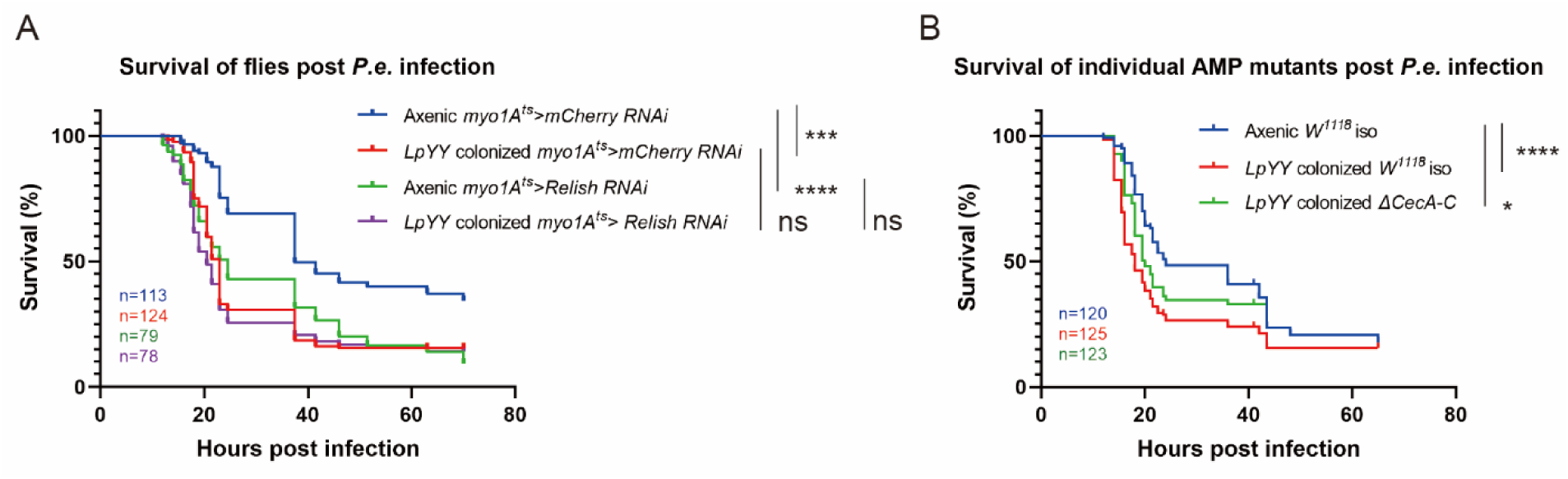
Reduced expression of antimicrobial peptides reduces the hyper susceptibility of *Lp*-colonized flies to *Pe* infection. (A) Survival curves of *Lactiplantibacillus* and axenic flies with gut-specific *Relish* downregulation (*myo1A^ts^*>*Relish RNAi*) upon *P. entomophila* infection. *myo1A^ts^*>*mCherry RNAi* was used as a control. The accumulated survival graph is shown (*n* = 2-3). (B) Survival curves of *Lactiplantibacillus-*colonized *w^1118^* iso flies or flies deleted with antimicrobial peptides *CecA-C* upon *P. entomophila* infection. The accumulated survival graph is shown (*n* = 3). Data analyzed with stratified Cox hazard ratios by R (A and B). ns, not significant; \**p* < 0.05, \*\**p* < 0.01, \*\*\**p* < 0.001, \*\*\*\**p* < 0.0001. For detailed sample sizes and statistical analyses, see Table S1.

**Figure S3.**
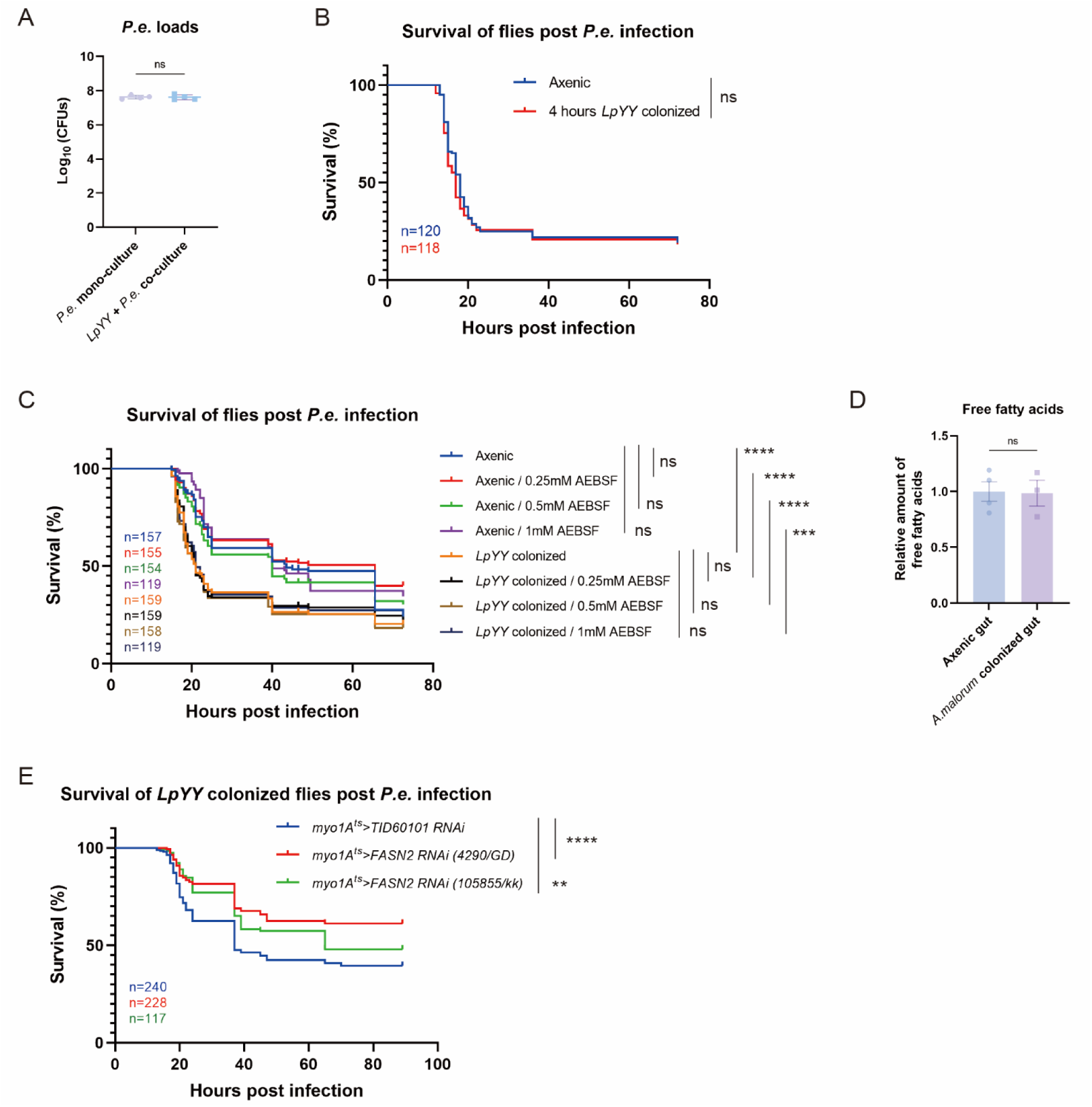
*Lp* facilitates *Pe* infection in flies by enhancing host fatty acid synthesis and not protease expression. (A) *Pe* loads (CFUs) at 24 hours mono- or co-culture with *LpYY* in BHI medium (*n* = 4). Bars show mean ± SEM. (B) Survival curves of axenic *w^1118^* iso flies and flies colonized with *Lactiplantibacillus* for 4 hours upon *P. entomophila* infection. The accumulated survival graph is shown (*n* = 3). (C) Survival curves of axenic *w^1118^* iso flies and *Lactiplantibacillus-*colonized flies pretreated with the protease inhibitor AEBSF upon *P. entomophila* infection. The accumulated survival graph is shown (*n* = 3). (D) Free fatty acid measurements from the guts of *Acetobacter*-colonized axenic *w^1118^* iso flies and axenic flies. The relative level of free fatty acids was normalized by setting the value from the axenic gut to 1. Each dot indicates a pool of 20 fly guts taken from different infection vials (*n* ≥ 3, pooled from 2 experiments). Bars show mean ± SEM. (E) Survival curves of *Lactiplantibacillus* flies with gut-specific downregulation of fatty acid synthase 2 (*FASN2*) upon *P. entomophila* infection and using the TID60101 RNAi line as a control (*myo1A^ts^>* TID60101 RNAi). The accumulated survival graph is shown (*n* = 4).

**Figure S4.**
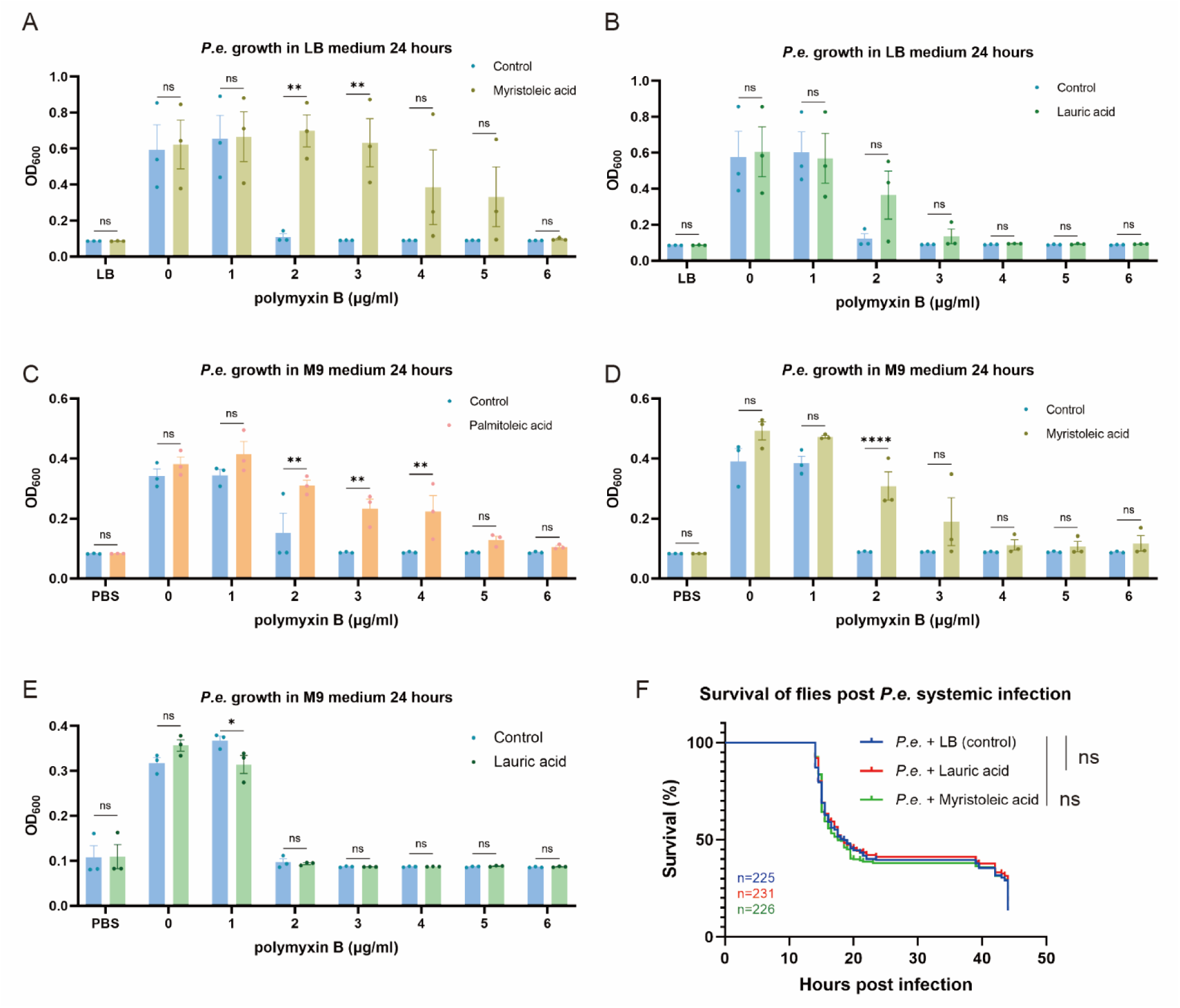
Myristoleic acid and lauric acid do not increase *Pe* virulence. (A and B) Growth of *P. entomophila* in LB medium with different concentrations of polymyxin B or with polymyxin B plus 0.3 mM myristoleic acid (A) or 0.3 mM lauric acid (B). Bars show mean ± SEM. (C, D and E) Growth of *P. entomophila* in M9 medium with different concentrations of polymyxin B or with polymyxin B plus 0.5 mM palmitoleic acid (C), 0.3 mM myristoleic acid (D) or 0.3 mM lauric acid (E). Bars show mean ± SEM. (F) Survival curves of axenic *w^1118^* iso flies systemically infected with *P. entomophila* by pricking or *P. entomophila* pre-incubated with 0.3 mM myristoleic acid 0.3 mM, palmitoleic acid, or 0.3 mM lauric acid for 3 hours. The accumulated survival graph is shown (*n* = 5). Data analyzed with an unpaired t -test (A, B, C, D, and E) and stratified Cox hazard ratios (F). ns, not significant; \**p* < 0.05, \*\**p* < 0.01, \*\*\**p* < 0.001, \*\*\*\**p* < 0.0001. For detailed sample sizes and statistical analyses, see Table S1.

## Notes

### Competing Interest Statement

The authors have declared no competing interest.

